# Assessing *in vivo* the impact of gene context on transcription through DNA supercoiling

**DOI:** 10.1101/2022.10.31.514473

**Authors:** Ihab Boulas, Lisa Bruno, Sylvie Rimsky, Olivier Espeli, Ivan Junier, Olivier Rivoire

## Abstract

Gene context can have significant impact on gene expression but is currently not integrated in quantitative models of gene regulation despite known biophysical principles and quantitative *in vitro* measurements. Conceptually, the simplest gene context consists of a single gene framed by two topological barriers, known as the twin transcriptional-loop model, which illustrates the interplay between transcription and DNA supercoiling. *In vivo*, DNA supercoiling is additionally modulated by topoisomerases, whose modus operandi remains to be quantified. Here, we bridge the gap between theory and *in vivo* properties by realizing in *Escherichia coli* the twin transcriptional-loop model and by measuring how gene expression varies with promoters and distances to the topological barriers. We find that gene expression depends on the distance to the upstream barrier but not to the downstream barrier, with a promoter-dependent intensity. We rationalize these findings with a first-principle biophysical model of DNA transcription. Our results are explained if TopoI and gyrase both act specifically, respectively upstream and downstream of the gene, with antagonistic effects of TopoI, which can repress initiation while facilitating elongation. Altogether, our work sets the foundations for a systematic and quantitative description of the impact of gene context on gene regulation.

## 1 Introduction

Gene regulation is most often studied through the lens of transcription factors, leading to its representation as regulatory networks where gene context – the relative location and orientation of genes along DNA – is abstracted away. This simplification has important limitations. It cannot explain, for instance, how reduced bacterial genomes with very few transcription factors generate intricate patterns of gene expression [1, 2, 3]. While multiple factors other than transcription factors may be invoked [4, 5, 6], the confrontation of transcriptional data with comparative genomics reveals that gene context plays a primary role, at least in bacteria [7]. Accordingly, the expression of a transcription reporter cassette depends strongly on its location along the *E. coli* chromosome [8]. Similarly, on a plasmid, the relative orientation of genes has a significant impact on their expression levels [9]. Experimental data also show that a given synthetic regulatory network can behave qualitatively differently in different genetic contexts. [10]. Genome organization is correspondingly found to be evolutionarily more conserved than transcription factor regulation in natural genomes [11]. Yet, how gene context affects gene expression remains poorly understood.

Gene context may impact gene expression in different ways. In bacteria, a simple but pervasive effect is transcriptional read-through, where the absence or the over-riding of terminators cause a downstream co-directional gene to be co-transcribed with an upstream gene [12]. RNA polymerases (RNAPs) may also interact physically, leading to different forms of transcriptional interference [13]. Additionally, the activity of different RNAPs may be coupled indirectly through mechanical perturbation of DNA. Supercoiling, the over- or under-winding of the double helix, is indeed known to affect and to be affected by transcription [14, 15, 16]: as an RNAP transcribes, it exerts a mechanical stress on DNA which causes the double helix to be under-wound upstream and over-wound down-stream of the gene [17]. This mechanical perturbation can propagate through distances of several kilo-bases [18] to affect neighboring or subsequent initiations [19] and elongations [15] of transcription. Finally, several proteins can impact transcription by modulating supercoiling. These include topoisomerases, which regulate DNA supercoiling [20], as well as nucleoid-associated proteins [21] which may form topological barriers and prevent the diffusion of supercoiling [22, 23].

Conceptually, the simplest situation where gene context can impact gene expression involves a single gene framed by two topological barriers that prevent the diffusion of DNA supercoiling (Fig. 1A). This defines the “twin transcriptional-loop model” introduced thirty five years ago by Liu and Wang to illustrate the interplay between transcription and supercoiling [17], with negative and positive DNA supercoiling generated upstream and, respectively, downstream of an elongating RNAP (Fig. 1A). This model is nowadays at the foundation of all theoretical studies of the impact of gene context on gene expression [24, 25, 9, 26, 27]. It is also central to multiple *in vitro* single-molecule experiments that have led to many insights on the translocation of RNAPs along DNA and on the activity of topoisomerases [28, 29, 30, 15]. As a result, mechanical and topological constraints generated during transcription are well understood at a quantitative level.

**Figure 1:**
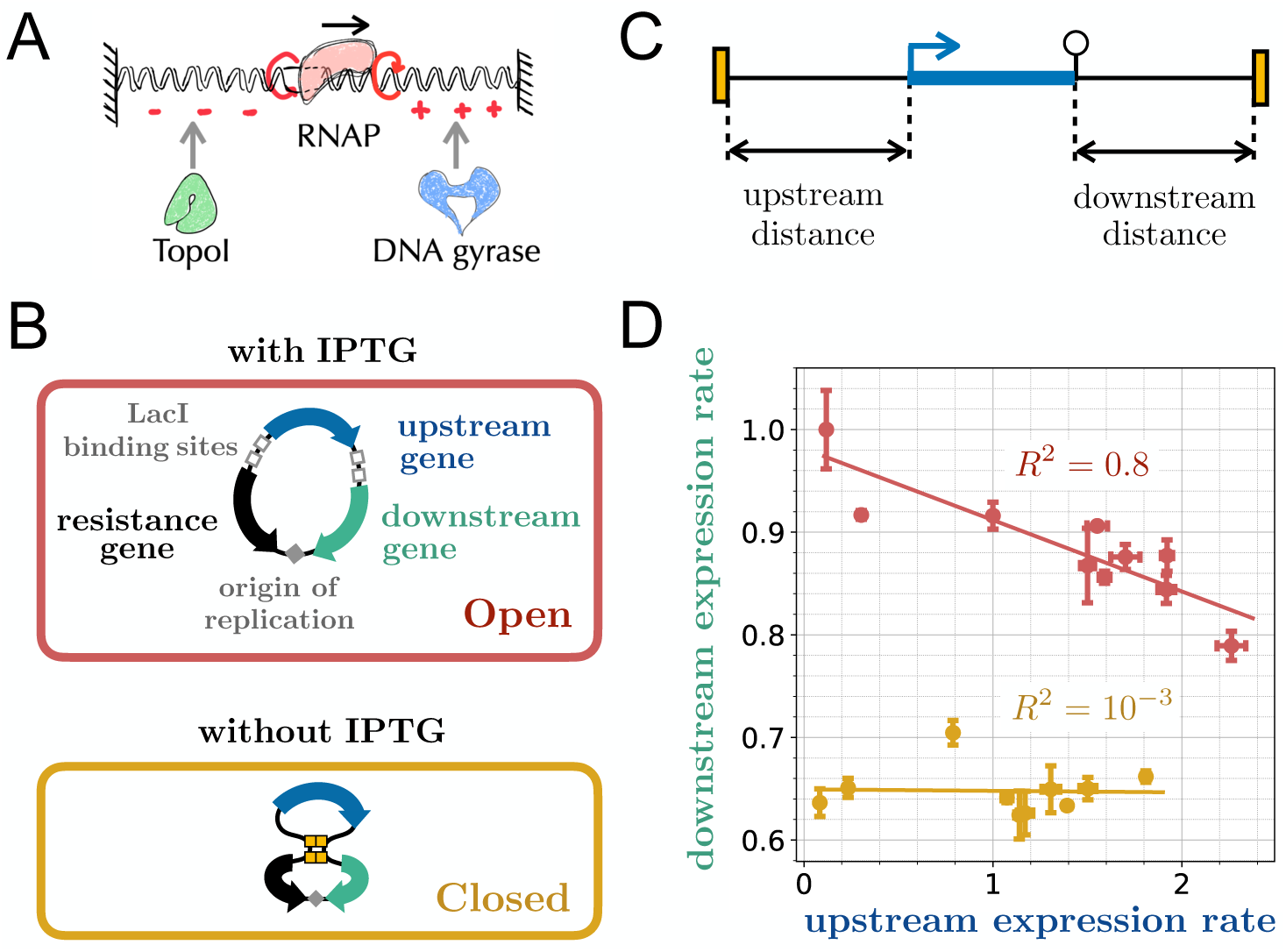
Conceptual and experimental models – **A.** The conceptual “twin transcriptional-loop model” consists of a single gene delimited by two barriers that prevent the diffusion of supercoiling [17]. A transcribing RNA polymerase generates negative supercoiling upstream and positive supercoiling downstream, which may eventually hinder further transcription due to torsional torques. In *E. coli*, just as in most bacteria, mainly two topoisomerases can resolve these constraints: TopoI, which relaxes negative supercoils, and DNA gyrase, which relaxes positive ones. **B.** We implemented this model on a plasmid with two genes coding for fluorescent proteins, here indicated as upstream and downstream genes, and an antibiotics resistance gene. The upstream gene is flanked by tandems of LacI binding sites. In absence of IPTG, LacI forms two loops between which supercoiling cannot diffuse [22, 23], thus insulating the upstream gene. **C.** We built several such systems that differ by the promoter sequence of the upstream gene and the downstream and upstream distances from the promoter or terminator to the boundaries, which are joined by LacI in the closed system. **D.** Expression rate of the downstream gene versus expression rate of the upstream gene for given distances but different promoters of the upstream gene (Tab. S1), measured either in the open (in red) or closed (in yellow) system. Downstream expression rates are normalized by their largest value and upstream expression rates by that of the promoter used downstream when placed upstream. While the expression rates of the downstream and upstream genes are negatively correlated when the system is open, they become uncorrelated when it is closed, consistent with their transcriptional insulation.

The application of the twin transcriptional-loop model to account for *in vivo* phenomena faces, however, two main difficulties. First, our quantitative understanding is limited with respect to the *in vivo* impact of topoisomerases on DNA supercoiling. While several topoisomerases are known to manipulate the topology of DNA, the two main topoisomerases implicated in transcription in *E. coli* are DNA gyrase, which removes positive supercoils, and TopoI, which removes negative supercoils [31]. Their *in vivo* activities are, however, not known quantitatively. For instance, high-throughput *in vitro* single-molecule assays suggest that the accumulation of positive supercoiling ahead of transcription and its transient release by gyrase produces transcriptional bursts [32] but whether this scenario explains the burst observed *in vivo* depends critically on whether gyrase is limiting *in vivo*, as it has been shown for instance for TopoI [33]. The issue is not only quantitative as the main mode of action of topoisomerases is also not clear: topoisomerases may indeed act either unspecifically or specifically, where specificity may involve DNA motifs [20], DNA mechanical states [34], or interactions with RNAPs [35, 36]. A second difficulty is the diversity of promoter sequences present in genomes, which are well known to differ not only in strength but in their response to DNA supercoiling [37]. These phenotypes cannot currently be predicted accurately from promoter sequences and generally conceal a diversity of underlying physical parameters, including binding, unbinding and initiation rates. As a consequence of these two difficulties, our conceptual and *in vitro* understanding of the interplay between transcription and DNA mechanics cannot presently be applied to a quantitative description of the *in vivo* impact of gene context on gene expression.

Here, we address these difficulties by implementing *in vivo* in *E. coli* different instances of the twin transcriptional-loop model with a single gene insulated from its neighbors (Fig. 1B). We realize this insulation using DNA bridging proteins that we place at varying distances to a range of different promoters (Fig. 1C) and use the data to constrain a first-principle biophysical model of gene transcription where the only free parameters are the mode and intensity of action of topoisomerases. The resulting theoretical model accounts quantitatively for our experimental results and further makes predictions on the mode of action of topoisomerases. Altogether, the combination of our experimental and theoretical models provides a critical missing link between conceptual models, *in vitro* measurements and *in vivo* phenomena, thus paving the way towards a quantitative understanding of the impact of gene contexts on gene expression.

## 2 Material & Methods

### 2.1 **Experimental methods**

#### Strains and plasmids

All measurements were carried out in the *E. coli* MG1655 background. The genetic constructs for the minimal system use the pSC101 origin of replication making it a low copy plasmid and Kanamycin resistant. The upstream gene encodes the fluorescent protein mCerulean ME, and the downstream gene the fluorescent protein mVenus ME. Their very similar sequences, comparable folding time and long life times allow for a straightforward comparison of their expression rates. The terminators B0014 and T1 follow mCerulean and mVenus, respectively (see Table S2 for their sequences). For the downstream gene, the Ribosome Binding Site (RBS) is always the same (Table S2) and the promoter is always pR – except for Fig. 1 where it is apFAB61 (Table S1). For the upstream gene, the RBS is always apFAB837 (Table S2) and the promoter sequences used can be found in Table S1. In Fig. 3, the weak, medium and strong promoters are apFAB45, apFAB67 and apFAB70, respectively. Each topological barrier is composed of two tandem lacO biding sites (the plasmid has therefore 4 lacO binding sites in total). The two barriers are also in a tandem orientation with one another. Their sequence differs slightly from that of [22] to avoid unnecessary repeats. Their sequences can be found in Table S1. The upstream and downstream distances to the barriers were obtained from the PCR of regions of the *λ* phage genome, which is unlikely to contain cryptic promoters [38]. For Figures S18 and S19, opB::kan, gyrBts::tet and parEts::tet alleles were introduced in MG1655 by P1 transduction [39].

#### Growth medium

All of the experiments were carried out in M9 minimal medium using the following recipe: 1X M9 Minimal Salts (from Sigma Aldrich); 0.4% Glucose; 1% Casaminoacids ; 2 mM MgSO_4_ and 0.1 mM CaCl_2_. In addition, Kanamycin was added at 50 *µ*g/mL. To obtain an open loop, 1mM of IPTG is added to the medium.

#### Data acquisition

Glycerol stocks were streaked on resistance agar plates. Single colonies were inoculated in 1 mL of minimal medium with antibiotics, within a 2 mL 96 deepwell plate. Cultures were grown overnight in a thermoblock, at 37^◦^ C and 1200 rpm. Cultures then underwent a 1:500 dilution in 1 mL of minimal medium with antibiotics, within a 2 mL 96 deepwell plate. An outgrowth was run in a thermoblock, at 37^◦^ C and 12,000 rpm, for 3h (to an OD_600_ 0.1). Cultures then underwent a 1:10,000 dilution and 100 *µ*L of these diluted cultures were aliquoted in a 96 well plates (with black walls and a clear flat bottom). 50 *µ*L of mineral oil was finally added to each well. Time series were acquired in a Tecan Spark Microplate Reader. No Humidity Cassette was used. Temperature was set at 37^◦^ C (at least 1h prior to the beginning of the acquisition). Shaking was set on double orbital with amplitude of 3 mm and frequency of 90 rpm. Time points were acquired every 25 minutes, over a total period of *∼* 20h. Three quantities were measured at each time point: absorbance (at 600 nm); mCerulean fluorescence (excitation at 430 nm and emission at 475 nm using a manual gain of 90) and mVenus fluorescence (excitation at 510 nm and emission at 550 using a manual gain of 70).

One-dimensional gel electrophoresis with chloroquine. Single colonies of *E. coli* MG1655 WT or GyrBts harboring the desired plasmids were inoculated in 1 mL of minimal medium with Kanamycin (50 *µ*g/mL). Cultures were grown overnight in an incubator at 30^◦^C, 180 rpm. For each strain, three flasks containing 50 mL of minimal medium with Kanamycin were inoculated with the overnight cell culture at 1:500 dilution. The cultures were grown respectively at 30, 33 and 37^◦^C with agitation (180 rpm) until OD= 0.2. Plasmid DNA molecules were purified using a commercially available purification kit (Monarch*Q*R Plasmid Miniprep Kit, NEB). The purified plasmids were run on a 0.8% agarose gel supplemented with 2.5 *µ*g/mL of chloroquine in 1xTBE buffer containing 2.5 *µ*g/mL of chloroquine at 25 V for 15 hours. The agarose gels were then washed in tap water three times during 30 minutes, and stained by SYBR Green.

### 2.2 **Inference of gene expression rates**

#### Preprocessing

First, the raw temporal data for the optical density and fluorescence is linearly interpolated over 750 points, from the ~ 50 raw data points (using the interp1d module from the SciPy library in Python). The interpolated data is then filtered using a 2^nd^ order polynomial (by a Savitzki-Golay filter using the savgol module from the SciPy library in Python using a window size of 101). The relative differences in gene expression rates are only weakly sensitive to the exact parameters used for the interpolation and filtering.

#### Computation of expression rates

The gene expression rate (as represented in Fig. 1) is computed as *α_F_* = (*dF_t_/dt*)*/*(*dN_t_/dt*), where *F_t_* is the fluorescence signal and *N_t_* is the optical density signal (see Figs. S11-S13 for a justification and Figs. S14-S16 for a comparison with other approaches). This computation allows for the identification of a region (between the background-dominated early phase and the entrance into stationary phase) during which gene expression rate is stable (Fig. S12). This region is identified automatically by minimizing the signal derivative over a temporal region of ~ 1h45. If slower growth (at 29^◦^ C instead of 37^◦^ C) is used, the duration of the stable signal region extends to 5h (Fig. S13). Gene expression rate is temporally averaged over this stable region.

#### Normalization of expression rates

The gene expression rate is obtained both for the upstream and downstream genes. They are independently compared in Fig. 1. For Figs. 2-3, the upstream gene expression rate is normalized by that of the downstream gene to remove copy number differences due to changes in plasmid size. This normalization is justified by the independence between the two genes demonstrated in Fig. 1. The relative gene expression rate (as represented in Fig. 2-3) is computed as 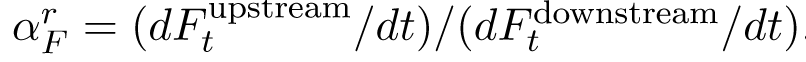. As for the simple gene expression computation, this computation yields a stable region of gene expression rate which is identified and averaged.

**Figure 2:**
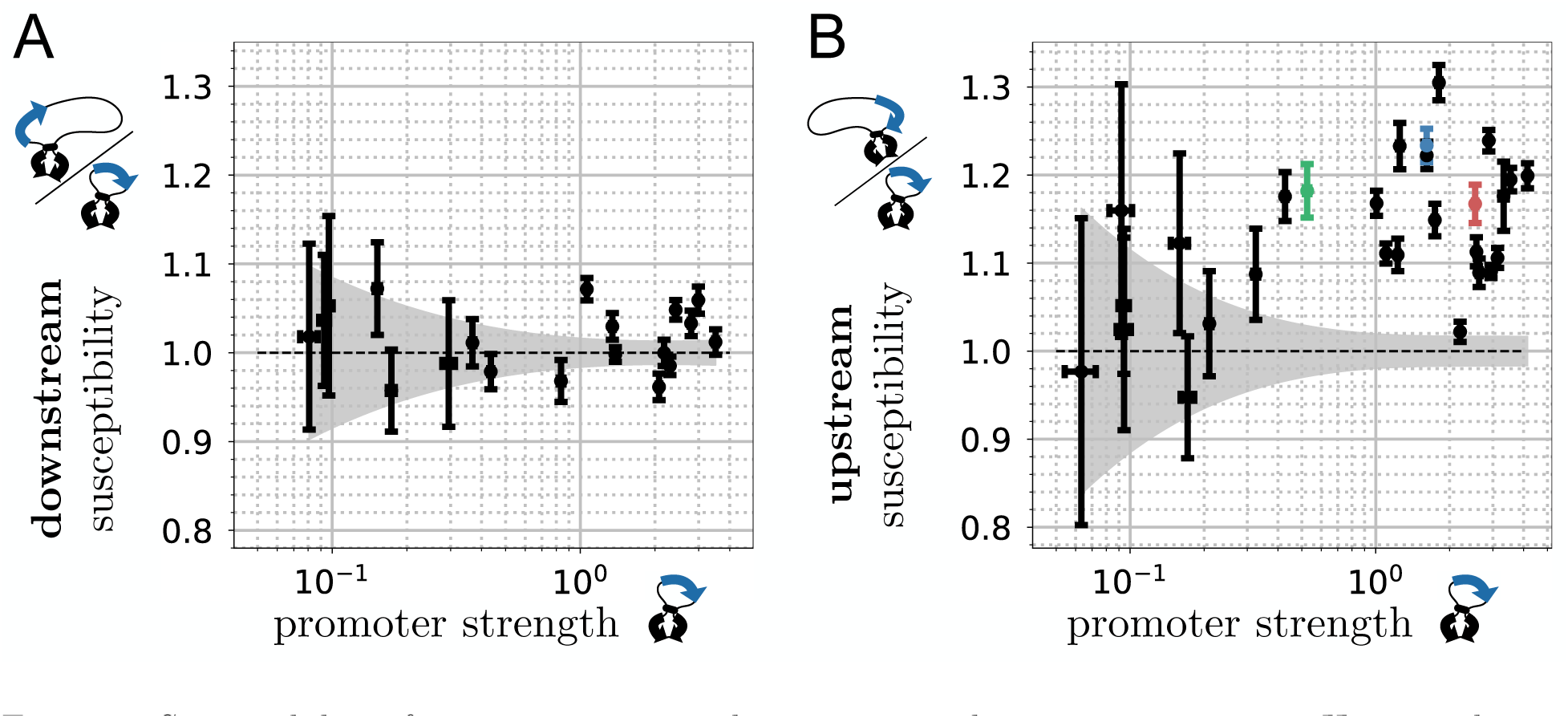
Susceptibility of gene expression to downstream and upstream contexts – Here we change the context of the insulated gene by introducing a 3 kb sequence either downstream or upstream and consider promoters of varying strengths. **A.** Susceptibility to downstream context versus promoter strength. The downstream susceptibility is defined as the ratio of the expression rate of the insulated gene with a long (3408 bp) distance to the downstream barrier over its expression rate with a short (320 bp) distance. Measurements involving weak promoters are less precise as indicated by the shaded area marking a deviation from unity by less than one standard deviation across replicate measurements (Methods). **B.** Susceptibility to upstream context versus promoter strength. The upstream susceptibility is defined as the ratio of the expression rate of the insulated gene with a long (3200 bp) distance to the upstream barrier over its expression rate with a short (250 bp) distance. In contrast with downstream susceptibility, it is significantly larger than one for all but one of the strong promoters. The three promoters marked in color are further studied in Fig. 3.

#### Susceptibilities

The relative expression rates are obtained for different gene contexts. The strength of a promoter is defined as the relative expression rate measured when both the upstream and downstream distances are short. The susceptibility to the upstream context is the ratio of relative expression rate in the long upstream context over that of in short upstream context, and conversely for the susceptibility to the downstream context. Note, here, that promoter strength should actually be measured in the absence of any context effect. This is nevertheless never the case in practice. We thus checked in our biophysical model that results and conclusions are identical when defining promoter strength from long distances.

#### Errorbars

For each data point, 4 to 8 replica (constituted of different colonies from a given glycerol stock) were made. Because the inferred gene expression rate comes from a temporal average, we computed the error associated to these replica as the standard error of the mean (their magnitude can be seen in the *x* and *y* axes of Fig. 1D for example). The propagation of error when ratios of average gene expression rates are considered (as seen in the *y* axis of Fig.2A-B) is approximated as

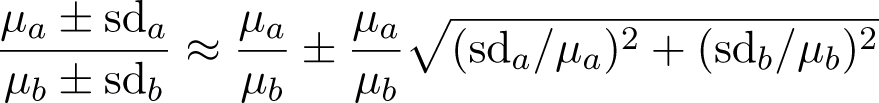

where *µ* stands for mean and sd for standard deviation. In Figs. 2A-B, the grey shadow is obtained by plotting the magnitude of the ratio errors and fitting a 3^rd^ degree polynomial to them. When statistical significance is computed, it is via the independent t-test.

### 2.3 **Biophysical model**

#### Definitions

The model considers a segment of DNA discretized at the single base level into sites with topological barriers at *x* = 0 and *x* = *L*, a TSS at *x* = *d* and a terminator at *x* = *d* + *L_g_*(Fig. 4). The TSS can be occupied by a non-elongating RNAP and sites *X*_1_*, …, X_N_* at *d X*_1_ *< < X_N_ < x_p_* + *L_g_* by a varying number *N* of elongating RNAPs, and we define *X*_0_ = 0. Binding can occur if the TSS is free, that is, if *N* = 0 or if *X*_1_ *d > £*_RNAP_, with *£*_RNAP_ the exclusion length of an elongating RNAP. Elongating RNAPs constitute topological barriers and supercoiling relaxes quickly with respect to the time scale of elongation to take an uniform value Σ*_i_* between *X_i_*_−1_ and *X_i_*. Σ*_i_* = *n*Lk*_i_/*(*X_i_ − X_i_*_−1_) 1, where Lk*_i_* is the corresponding linking number and *n* = 10.5 the number of base pairs per DNA helix. We also define Lk*_N_*_+1_, the linking number of the DNA downstream of the most downstream RNAP and Lk_0_ the linking number of the domain when no elongating RNAP is present, and *X_N_*_+1_ = *L* such that Σ*_N_*_+1_ = *n*Lk*_N_*_+1_*/*(*X_N_*_+1_ *X_N_*) 1.

#### Simulations

The dynamics are simulated in discrete time with time unit *τ*_0_ = *v*^−1^ where *v_m_* is the RNAP translocation speed in bp.s^−1^. Starting from *N* = 0, a simulation run consisted in performing the following updates, with *T* the total time of the simulation (in practice, we use a Gillespie algorithm to speed up the simulations):

(1) A new RNAP binds at *d* with probability *k_b_τ*_0_Θ(*X*_1_ *x_p_ £*_RNAP_)(1 − *δ_p_*) where Θ represents the Heaviside function and *δ_p_* = 1 if the promoter is bound by an RNAP, 0 otherwise.
(2) An RNAP bound at *d* is considered to form a closed complex with DNA. We then consider the transition to the open complex to occur with probability *k_o_τ*_0_Θ(*σ_o_* − Σ_1_) where Σ_1_ is the supercoiling density behind the last elongating RNAP if there is one (*N >* 0) and Σ_1_ *< σ_o_* otherwise (*N* = 0).
(3) The initiation of elongation occurs once the open complex is formed with probability *k_e_τ*_0_ (or 1 in the case where *k_e_ >* 1*/τ*_0_). The newly elongating RNAP is labeled *i* = 1 and the following updates are made: *N N* +1, *X*_1_ *d*, Σ_1_ Σ_1_, Lk_1_ (1+Σ_1_)*d/n* and, for *i >* 1, *X_i_ X_i_*_−1_, Σ*_i_* Σ*_i_*_−1_ and Lk*_i_* (1 + Σ*_i_*_−1_)(*X_i_ X_i_*_−1_)*/n* except Lk_2_, which is updated as Lk_2_ (1 + Σ_1_)(*X*_1_ − *d*)*/n*.
(4) In the presence of at least one elongating RNAP (i.e., *N* ≥ 1), the linking numbers of the upstream and downstream part of the system, Lk_1_ and Lk*_N_*_+1_, are updated to account for the actions of TopoI and gyrase. For the non-specific activities, we have Lk_1_ *←* Lk_1_ + *A_T_ −* 2*A_G_* and Lk*_N_*_+1_ *←* Lk*_N_*_+1_ + *A*^t^ *−* 2*A_G_* where, on the one hand, *A_T_* and *A*^t^′ are random variables associated with TopoI activity and drawn from a Poisson distribution with mean *τ λ*^Topo^_*ns*_*X* (0 if 0 *ns* 1 Σ_1_ *> −*0.05 [40]) and, on the other hand, *A_G_* and *A*^t^ are random variables associated with gyrase activity and drawn from a Poisson distribution with mean *τ λ*^Gyr^_*ns*_(*L – X*)*/*2 (0 if Σ *< σ* to prevent supercoiling from drifting away). For the specific activities, we have: Lk_1_ *←* Lk_1_ + 1 and Lk *←* Lk *−* 2 with respective probabilities *τ*_0_Λ^Topo^_*ns*_ (0 if Σ *< −*0.05) and *τ*_0_Λ^Gyr^*_s_*/2. In the absence of any elongating RNAP, only non-specific activities are considered, and we have Lk *←* Lk + *A −* 2*A* with mean of the Poisson random variables *A* and *A* given by *τ_0_ λ*^Topo^_*ns*_*L* and *τ_0_ λ*^Gyr^_*ns*_*L*/2, respectively.
(5) Each RNAP *i*, whose order is taken randomly (asynchronous update), moves forwards *X_i_ X_i_*+1 with probability Θ(Σ*_i_ − σ_s_*) – to avoid artifacts from the discrete nature of the dynamics, we do not consider any exclusion effect between two consecutive elongating RNAPs, supercoiling constrains already preventing RNAPs to pass each other). Following this update, the linking numbers are unchanged but the densities of supercoiling Σ*_i_* and Σ*_i_*_+1_ are updated to account for the new distances *X_i_ X_i_*_−1_ and *X_i_*_+1_ *X_i_*.
(6) Any RNAP reaching the terminator at *d* + *L_g_* is removed (*N N* 1) and contributes to increase the number of transcripts by one: *M M* + 1.
(7) *T T* + *τ*_0_.

#### Computation of transcription rates

Using this framework, we estimate the transcription rate *ρ*(*d*) of a wide range of promoters (Table 1) at various upstream distances *d* and downstream distance *L L_g_*. This rate is obtained as the number of transcripts *M* obtained per total time *T*, *ρ* = *M/T*. Just as in the experiments, upstream susceptibilities are computed using *ρ*(*d*)*/ρ*(*d* = 250 bp). *M* is taken sufficiently large to get an unbiased estimation of the stationary transcription rate: *M* = 10^4^ when testing the full range of promoters (Fig. 5A) and *M* = 10^5^ when analyzing in more detail specific promoters (Fig. 5B).

**Table 1:** Model parameters – Our biophysical model involves three types of parameters: **A.** system parameters whose value is either known from the literature or fixed by experimental design; **B.** parameters characterizing each promoter, which are generally unknown and for which we consider a range of values; **C.** unknown parameters related to the *in vivo* activity of topoisomerases, which we estimate using our experimental results. SC stands for supercoiling, OC for open complex, Topo for topoisomerase I and Gyr for gyrase.

| **A. Known parameters** |  |  |
| --- | --- | --- |
| $L_g$ | gene length | 900 bp |
| $L$ | distance between the 2 barriers | varying |
| $d$ | distance to upstream barrier | varying |
| $n$ | number of bp per B-DNA helix | 10.5 bp |
| $\ell_{\text{RNAP}}$ | RNAP exclusion length | 30 bp |
| $v_m$ | elongation speed | 25 bp.s<sup>-1</sup> |
| $\sigma_s$ | SC threshold for elongation | -0.062 |
| $\lambda_{ns}^{\text{Gyr}}$ | gyrase non-specific activity | -10<sup>-4</sup> Lk.bp<sup>-1</sup>.s<sup>-1</sup> |
| **B. Expected range of promoter parameters** |  |  |
| $k_b$ | binding rate | $\in [0.01, 10.24]\text{s}^{-1}$ |
| $k_o$ | basal rate for OC formation | $\in [0.01, 10.24]\text{s}^{-1}$ |
| $\sigma_o$ | SC threshold for OC formation | $\in [-0.05, 0]$ |
| $k_e$ | escape rate | $\in [0.01, 10.24]\text{s}^{-1}$ |
| **C. Unknown topoisomerase parameters** |  |  |
| $\lambda_{ns}^{\text{Topo}}$ | TopoI non-specific activity | |
| $\Lambda_s^{\text{Gyr}}$ | Gyr specific downstream activity | |
| $\Lambda_s^{\text{Topo}}$ | TopoI specific upstream activity | |

In Fig. 5A and Fig. 6C-D, we tested 2646 combinations of values of *k_b_*, *k_o_* and *σ_o_* where the fre-quencies were taken in *{*0.01*×^√^*2*^i^}* and *σ* in *{−*0.05*, −*0.04*, −*0.03*, −*0.02*, −*0.01, 0*}* (*k τ* = 1 *o e* 0 in these figures). In Fig. 5B, the following parameters are used for the weak promoter: *k_b_* = 0.64.s^−1^, *k_o_* = 0.04.s^−1^ and *σ_o_* = − 0.04; medium promoter: *k_b_* = 0.453.s^−1^, *k_o_* = 0.32.s^−1^ and *σ_o_* = − 0.05; strong promoter: *k_b_* = 0.32.s^−1^, *k_o_* = 3.62.s^−1^ and *σ_o_* = − 0.04. *k_e_τ*_0_ = 1 for the three promoters (immediate escape).

#### Stalling torque

The value of the stalling torque *σ_s_* = − 0.062 is based on the relationship *σ_s_* = Γ*_s_/A* [41], which holds for both super-structured DNA and unstructured DNA. Γ*_s_* = 18.5 pN.nm [42] is the RNAP stalling torque and *A* = 300 pN is chosen to be intermediate between the value estimated from single-molecule experiments for super-structured DNA (200 pN) and unstructured DNA (400 pN) [41].

#### OC formation rate

For the OC formation rate, we consider that the corresponding free energy barrier is reduced by DNA supercoiling independently of the promoter sequence [43]. In this context, the rate can be written as *k_o_* exp[*β*(Δ*G_P_* + Δ*G_σ_*)], where Δ*G_P_ >* 0 reflects sequence effect of the promoter and where Δ*G_σ_* reflects mechanical properties of DNA under supercoiling *σ*. *β*^−1^ = *k_B_T* is the energy unit, with *k_B_* the Boltzmann constant and *T* the temperature. We can then compare the rate *k_o_* exp[*β*(Δ*G_P_*)] in absence of supercoiling and the rate *k_o_*when Δ*G_σ_*compensates Δ*G_P_*. Δ*G_P_* being in general large with respect to *k_B_T* [44], we have *k_o_* exp[*β*(Δ*G_P_*)] *k_o_*. In accord with the sharp dependence of transcription rates as a function of *σ* [45], we then consider the compensation of Δ*G_P_* by Δ*G_σ_* to occur abruptly at a threshold value *σ_o_* that reflects Δ*G_P_*, i.e., *σ_o_* is promoter dependent. This eventually leads us to use *k_o_*Θ(*σ_o_ σ*) for the simplest form of the OC formation rate.

## 3 Results

### 3.1 An ***in vivo*** twin transcriptional-loop model

To design a genetic system where the transcription of one gene is insulated from the transcription of any other gene, we built on previous *in vitro* results showing how a pair of a tandem of protein binding sites (here *lacO* bound by LacI) can form topologically insulated loops that prevent the propagation of DNA supercoiling from one loop to the other [22, 23]. We introduced such binding sites on a plasmid comprising two co-directional genes separated by a strong terminator in addition to a resistance gene (Fig. 1B). The upstream fluorescent gene is placed in one loop to represent the insulated gene while the downstream fluorescent gene is placed with the resistance gene in the other loop.

The open system displays an interaction between the two fluorescent genes that illustrates the puzzling impact that gene context may have on gene expression: the activity of the downstream gene decreases linearly by up to 20% upon increasing the activity of the upstream gene by changing its promoter sequence (Fig. 1D). The simplest effect, transcriptional read-through, is inconsistent with the data, as it predicts the activity of the downstream gene to increase, not to decrease. Transcriptional interference, while predicting the downstream gene activity to decrease, also predicts that the upstream gene needs to be at least as expressed as the downstream gene to significantly affect it [46] while we observe that considerably weaker promoters have a significant impact on stronger ones (Fig. 1D). Other effects might then be hypothesized as for instance a repression of its initiation due to an excess of positive supercoiling generated by the upstream gene [24]. However, predicting the behavior of this three-gene system requires, first, to understand and quantify the mechanisms at play in the simpler, yet as we shall see already very rich case of a single insulated gene. This single-gene system, which is obtained by closing the loops, is an instance of the twin transcriptional-loop model and we verify that it effectively decouples the expression of the downstream gene from that of the upstream one (Fig. 1D).

### 3.2 **Experimental results**

#### 3.2.1 **Downstream versus upstream context**

We first study how transcription-induced supercoiling impacts gene expression *in vivo* by changing the upstream or downstream distances between the gene and the topological barriers (Fig. 1C). An increased distance is indeed expected to provide both more DNA to buffer the accumulation of supercoiling and more binding sites for supercoiling-managing topoisomerases (i.e., TopoI and gyrase) to relax this accumulation (Fig. 1A). If the accumulation of supercoiling has an impact on transcription, increasing these distances should therefore modify gene expression levels.

We thus designed different systems where we varied the promoter of the insulated gene (Table S1) and the distance either to the upstream barrier or to the downstream barrier, using promoter-free regions of the *λ* phage genome. We chose phage sequences because they have been thoroughly studied, and we verified that our results are reproduced with different sequences (Fig. S9). Our measurements were made independent of plasmid copy number (and therefore indirect plasmid size effects) as well as extrinsic factors of variability [47] by normalizing the gene expression rate of the insulated gene with that of a control gene located in the other topologically insulated loop (the “downstream gene” of Fig. 1B), thus defining a relative expression rate (Methods). To assess the sensitivity of gene expression to its downstream (or upstream) context, we compare this relative expression rate in a system with a long distance to the downstream (or upstream) barrier to that in a system with short distances to the two barriers. We call *downstream (or upstream) susceptibility* the ratio of the two rates. We then study these susceptibilities as a function of the promoter strength, which we take to be the relative expression rate measured when both the upstream and downstream distances are short (Methods). To this end, we selected several natural or synthetic promoters that cover a large range of expression strengths and are not known to be controlled by endogenous transcription factors [48] (Table S1).

When modifying the downstream distance between the stop codon and the barrier from 320 to 3408 base pairs (bp), we find that gene expression does not vary significantly, irrespectively of the promoter (Fig. 2A). In contrast, when modifying the upstream distance from 250 to 3205 bp between the transcription start site (TSS) and the barrier, we find gene expression to increase by up to 30% (Fig. 2B). For strong promoters (with strength at least 10 times larger than the smallest reported one), this gene expression amplification is statistically significant in all but one case. For weaker promoters, the measurements are less precise and, similarly to the downstream context, we find no evidence of susceptible promoters.

#### 3.2.2 **Dependence on promoter sequence**

The susceptibility to upstream context is not straightforwardly related to the promoter strength: the largest effect is obtained for a promoter whose promoter strength is two-fold smaller than the largest reported one and one of the strongest promoters is not susceptible at all (Fig. 2B). Can we rationalize this variability in terms of promoter sequence?

Two factors contribute to promoter strength: binding and initiation. Initiation can be further divided into two steps [49]: the formation of the open complex (OC), which involves a promoter-bound RNAP and a ~ 12 bp denatured DNA, and promoter escape. The formation of the OC has long been known to be sensitive to supercoiling [50]. Recent work [45] suggests this sensitivity to be primarily modulated by the GC content of a 6 bp long region preceding the start codon and, hence, located inside the so-called discriminator, i.e., the sequence downstream of the -10 hexamer [51, 52]. Here, however, we do not observe any significant correlation between the GC content of this region and the upstream susceptibility (Figs. S1-S2).

#### 3.2.3 **Dependence on upstream distance**

Is there an experimental parameter with a systematic impact on upstream susceptibility? Or might the variability of upstream susceptibilities from promoter to promoter conceal an uncontrolled source of variability? An answer is provided by analyzing in more depth how the expression of three specific promoters with different promoter strengths – referred in the sequel as “weak”, “medium” and “strong” – depends on the distance to the upstream barrier. In contrast to its intricate dependence on promoter sequence, upstream susceptibility indeed appears to have a simpler, monotonous dependence on the upstream distance (Fig. 3). More specifically, for the three investigated promoters, the susceptibility increases sub-linearly up to distances of the order of 1 kb, beyond which we observe two behaviors with no obvious relationship with promoter strength: on the one hand, the upstream susceptibility of the weak and strong promoters saturate at an amplification of approximately 20% while, on the other hand, that of the medium promoter keeps increasing roughly logarithmically.

**Figure 3:**
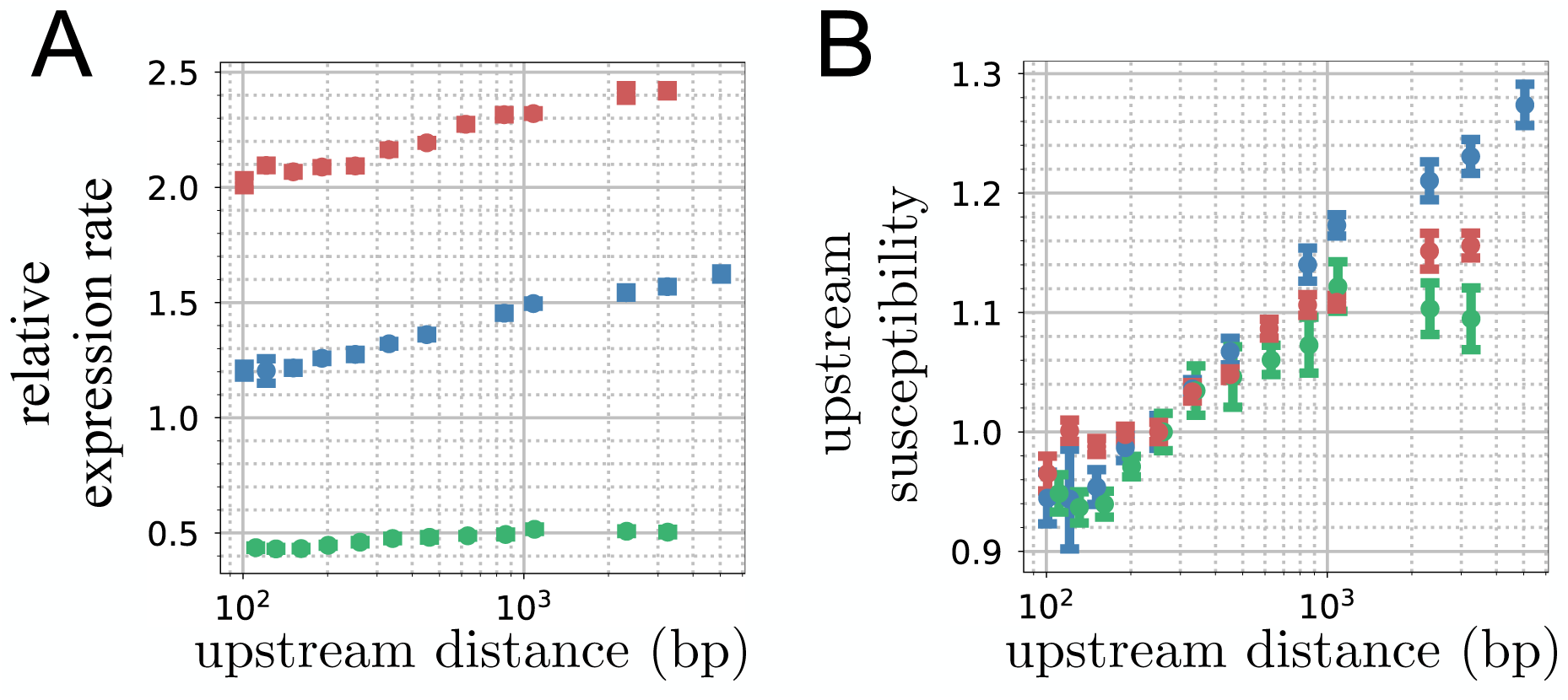
Dependence on distance to the upstream barrier – **A.** Relative expression rate when varying the distance to the upstream barrier for the three promoters marked in color in Fig. 2B – each point corresponds to an independent measurement. The downstream distance is here 520 bp, while it is 320 bp in Fig. 2B, explaining slight differences of upstream distance at 3205 bp. **B.** Susceptibility to upstream context for the same three promoters. The weak (green) and strong (red) promoters are found to have similar upstream susceptibilities despite having respectively a higher and lower promoter strength than the medium (blue) promoter.

We also examined the temperature dependence of the medium promoter, finding that its expression levels, but not its susceptibility to upstream context, depend on temperature (Fig. S3).

### 3.3 An *in silico* twin transcriptional-loop model

Can we build a first-principle biophysical model of the *in vivo* twin transcriptional-loop model that accounts for the different experimental results, namely (i) a susceptibility to upstream context but not to downstream context (Fig. 2), (ii) the dependence of the susceptibility to the distance to the upstream barrier (Fig. 3) and (iii) the non-trivial relationships between the upstream susceptibility and promoter strength (Figs. 2B and 3B)?

#### 3.3.1 **Modeling transcription**

To tackle this problem, we first built a minimal biophysical model of transcription by considering five major stages: RNAP binding to the promoter, formation of the OC, promoter escape, RNAP elongation and transcription termination (Fig. 4). Except termination, which is considered to occur immediately when an RNAP reaches the end of the gene, each of these stages is modeled as a stochastic process with a corresponding rate, with OC formation and RNAP elongation being the only processes sensitive to supercoiling (see below). We further constrain binding to occur only when the promoter is free, i.e., no other RNAP is present within *£*_RNAP_ = 30 bp [53]. As in previous quantitative models [24, 25, 26, 27] and consistent with *in vivo* experiments [54], we treat elongating RNAPs as topological barriers and assume DNA supercoiling to relax quickly relative to other time scales [23] so that the supercoiling density is uniform between successive topological barriers (Methods).

**Figure 4:**
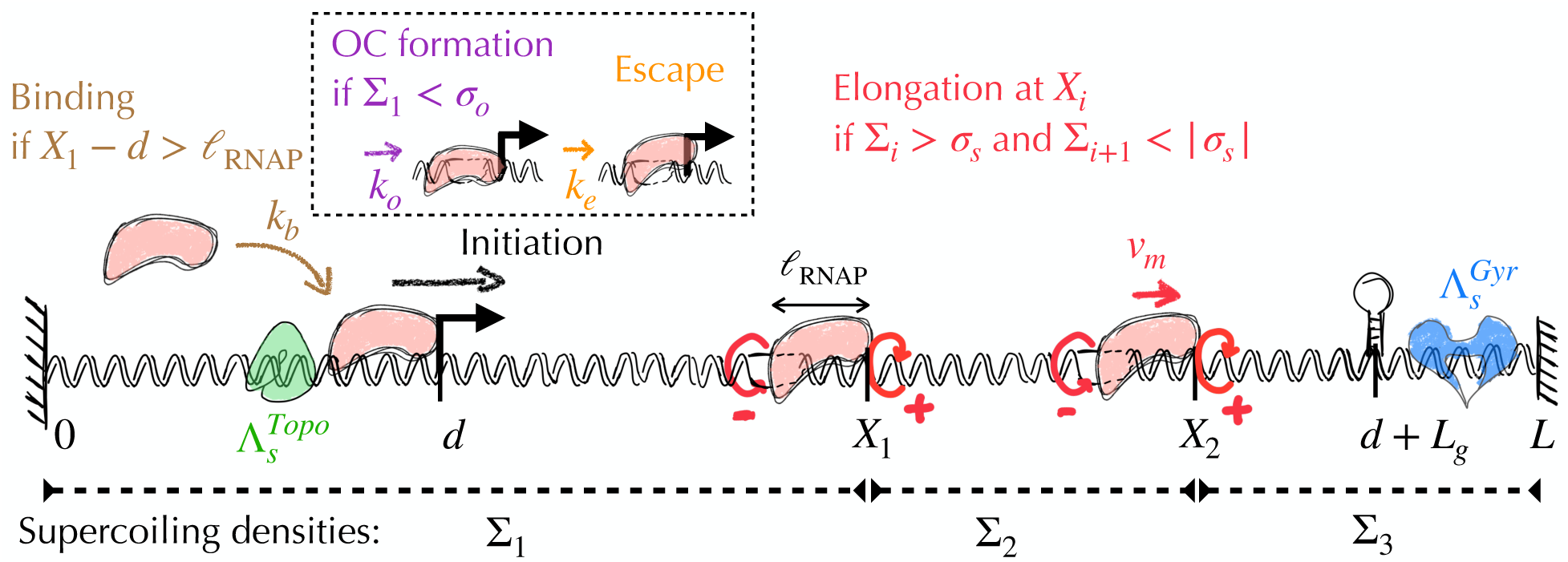
Schematic representation of our biophysical model of transcription under topological constraints – Transcription includes promoter binding by an RNAP, initiation of elongation which is divided into OC formation and promoter escape, elongation and termination. Elongating RNAPs behave as topological barriers and generate negative supercoils upstream (clockwise red arrows) and positive supercoils downstream (counterclockwise red arrows). The gene is embedded in a domain of length *L* that is topologically constrained at its extremities. If *N* RNAPs are elongating (here *N* = 2), *N* +1 independent topological domains are present whose supercoiling densities are denoted by Σ*_i_* (*i* = 1*, .., N* + 1). We further indicate the specific action of TopoI (green shape) and gyrase (blue shape) at the extremities of the gene. In addition, TopoI and gyrase may act non-specifically anywhere along the segment.

#### 3.3.2 **Known parameters**

Elongation involves fixed parameters known from single-molecule measurements (Table 1). Namely, RNAP translocation speed has been shown to be a sigmoid function of DNA supercoiling density [55]. Here, we consider a simple binary approximation of this dependence and assume elongating RNAPs to translocate at full speed *v_m_* provided the upstream (downstream) supercoiling densities are above (below) a supercoiling threshold *σ_s_* (*σ_s_*). Below *σ_s_* (above *σ_s_*), RNAPs remain immobile. *σ_s_* thus reflects RNAP stalling as a consequence of the large torque exerted by supercoiled DNA [55] and we take *σ_s_* = − 0.062 (Methods). For the elongation speed, we take *v_m_* = 25 bp.s^−1^, a value reported both in single-molecule *in vitro* experiments [55] and in *E. coli* growing in minimal medium [56], as used in our experiments.

#### 3.3.3 **Range of promoter-dependent parameters**

Binding, OC formation and promoter escape provide a coarse-grained decomposition of the multiple steps of transcription initiation [49, 44]. Kinetic details of each of these stages depend on promoter sequence, but the relationship remains poorly understood [44]. Here, to reflect the diversity of promoter sequences, we consider binding and escape to respectively occur at rates *k_b_* and *k_e_* with values uniformly sampled in the range [0.01, 10.24] s^−1^ [57, 44] (Table 1, Methods). For simplicity, we do not explicitly consider promoter unbinding, which we subsume in *k_b_*, nor abortive initiations, which we subsume in *k_e_*.

Next, OC formation involves DNA denaturation. Without DNA supercoiling, kinetics of this denaturation is slow, i.e., the corresponding free energy barrier is high and reflects promoter sequence [44]. DNA supercoiling reduces this barrier, mostly independently of the promoter sequence [43]. Considering that transcription initiation has a sharp sigmoid-like dependence on supercoiling [58] with a negligible rate above a certain threshold, the simplest description of the OC formation is to assume a non-zero rate *k_o_* only if the promoter supercoiling is below a threshold *σ_o_*(Methods). Here, we take *k_o_* in the same range as *k_b_* and *k_e_* ([0.01, 10.24] s^−1^) and *σ_o_* in the range [−0.05, 0] [45].

Finally, we found *k_b_*and *k_e_*to have very similar effects on the results (Fig. S4). For the sake of simplicity, we thus consider in the sequel that promoter escape is immediate once OC is formed (*k_e_* = *∞*) and discuss only the effect of *k_b_*.

#### 3.3.4 **Introducing topoisomerase activity**

In presence of topological barriers, transcription-generated DNA supercoils may generate strong variations of DNA supercoiling – all the stronger that barriers are closer – which need to be relaxed for transcription to proceed. We thus introduce in our model the stochastic action of TopoI, which removes negative supercoils, and of DNA gyrase, which removes positive ones. We assume that TopoI is active only when the supercoiling is below 0.05, as reported *in vivo* [40], and that gyrase is active only above *σ_s_* to prevent supercoiling from drifting away. Importantly, the *in vivo* modus operandi of topoisomerases remains poorly understood, with distinct scenarios being discussed in the literature (see e.g. [32, 33]). To be comprehensive, we thus consider two non-exclusive scenarios by which each of the two topoisomerases may relax DNA supercoiling.

On the one hand, topoisomerases may act non-specifically at any site (except, to simplify the handling of volume exclusion between DNA enzymes, between two elongating RNAPs). In this case, the corresponding removal rate of supercoils, *λ*^Topo^_*ns*_ and *λ*^Gyr^_*ns*_, are in units of Lk (linking number) per second and per base-pair, meaning that the non-specific activity of topoisomerases depends on the length of the corresponding topological domain. Based on *in vitro* measurement of activity and *in vivo* measurements of the density of active gyrases along DNA [59], we consider *λ*^Gyr^_*ns*_ = *−*10^−4^ Lk.bp^−1^.s^−1^. No corresponding measurement is available for *λ*^Topo^_*ns*_ and we therefore *ns ns* estimate below an upper bound value using our experimental results.

On the other hand, TopoI and gyrase may act specifically, i.e., at a precise location along the transcription process. In this case, the removal rates of supercoils, Λ^Topo^_*s*_ and Λ^Gyr^_*s*_, are in units of Lk per second, meaning that the specific activities of TopoI and gyrase do not depend on DNA length. Here, in agreement with the reported systematic localization of TopoI at the promoter of genes in various bacteria including *E. coli* [60, 61, 36], we consider the possibility for TopoI to act specifically upstream of transcribing RNAPs. In agreement with gyrase resolving the accumulation of positive supercoiling extremely efficiently [62] and having a biased distribution along bacterial genomes that reflects transcription activity [60, 34], we also consider the possibility for gyrase to act specifically downstream of transcribing RNAPs. As no *in vivo* measurement is available for Λ^Topo^_*s*_ and Λ^Gyr^_*s*_, we therefore use our experimental results to delineate possible values.

#### 3.3.5 **Simulations**

To implement the transcriptional-loop model, we embed a gene of fixed size *L_g_* = 900 bp in a larger domain of size *L* with the extremities *x* = 0 and *x* = *L* defining topological barriers (Methods, Fig. 4). The transcription start site is located at *x* = *d* such that the upstream and downstream distances are given by *d* and *L L_g_*, respectively. Simulations of the transcription process implement the stochastic dynamics of RNAP binding, OC formation, promoter escape, elongation and topoisomerase activities using a discrete-time approach. Transcription rates are measured in a stationary regime by computing the number of transcripts produced per unit of time. Susceptibilities are measured as in experiments by computing the ratio of transcription rates obtained at two different distances (Methods).

### 3.4 **Modeling results**

#### 3.4.1 **Parametrizing topoisomerase activity**

First, as the upstream distance increases and in absence of specific activity of TopoI, the non-specific activity of TopoI must increasingly contribute to the upstream susceptibility up to a characteristic distance on the order of *v_m_/*(*nλ*^Topo^_*ns*_) where the susceptibility saturates (Fig. S5; *n* = 10.5 is the number of base pairs per DNA helix). The absence of saturation for the medium promoter up to at least *d* max = 5 kb in Fig. 3B thus suggests *λ*^Topo^_*ns*_ to be smaller than *v_m_ /*(*nd*_max_) *∼* 5.10^−4^ Lk.bp^−1^.s^−1^. In the following we consider *λ*^Topo^_*ns*_ = 10^−4^ Lk.bp^−1^.s^−1^, identical to the known −λ^Gyr^_*ns*_

Second, in absence of any specific activity of either TopoI or gyrase, every stalling of an RNAP would last of the order of 10 to 1000 s. These correspond to the typical times for TopoI and gyrase to act through the non-specific mechanism, which are respectively given by (*λ*^Topo^_*ns*_*d*)^−1^ and (*λ*^Gyr^_*ns*_(*L − L*))^−1^. The removal rate of any promoter, including the strongest ones, would therefore be very low, considering *L_g_/n* 85 stalling events. This demonstrates the necessity to consider a specific activity for both TopoI and gyrase, respectively upstream and downstream the gene. We therefore tested a wide range of values of Λ^Topo,Gyr^_*ns*_ (Fig. S6) and assessed the capacity of the model to reproduce two properties of the dependence of the upstream susceptibility on promoter strength displayed in Figure 2B (where the largest distance is fixed to *d* = 3205 bp): a maximum susceptibility at *∼* 1.3 and the susceptibility of the strongest promoters lying between 1.1 and 1.2. The combination Λ^Gyr^_*s*_ = *−*2.5 Lk.s^−1^ and Λ^Topo^_*s*_ = 1.4 Lk.s^−1^ fulfills these requirements. We retain here these values but note that they are not the only one compatible with our results (Fig. S6). More generally, we find that Λ^Gyr^_*s*_ should be *≤ −*2 Lk.s^−1^ while the corresponding Λ^Topo^_*s*_ should lie between 1 and 2 Lk.s^−1^ (Fig. S6). Interestingly, *in vitro* single-molecule experiments have reported a similar value of removal rate of supercoils by TopoI, i.e., 1 Lk.s^−1^ [63]. In addition, our inference that Λ^Topo^_*s*_ *< |*Λ^Gyr^_*s*_*|* and Λ^Topo^_*s*_ *< v /n ∼* 2.4 Lk.s^−1^ is consistent with recent *in vivo* results showing that TopoI is limiting for transcription in *E. coli* [33].

#### 3.4.2 **Capturing promoter variability**

Given the above parameters, we can now study how the upstream susceptibility varies both from promoter to promoter and with respect to the distances to the topological barriers. First, we verify the absence of downstream effects (Fig. S7). This can be understood as a result of the limited impact of downstream barriers on elongation, primarily due to the low unspecific activity of DNA gyrase. Furthermore, experimental measurements and simulations are performed in a stationary regime where the average time between two transcript productions reflects initiation times rather than elongation times. In our model, for elongation to affect these initiation times, the most upstream RNAP must stall and block access of the promoter to new RNAPs, which is theoretically possible. However, this effect would only become apparent for extremely low values of the specific activities of topoisomerases, resulting in non-physiological elongation times.

Second, and consistent with Fig. 2B, we verify in Fig. 5A that the upstream susceptibility is not a simple function of promoter strength. More precisely, we obtain an overall shape of the distribution of susceptibilities very similar to experimental results where most of the weakest promoters are not susceptible and most of the strong promoters have a susceptibility above 1.1, with a large variability among strong promoters. The correspondence of the maximal value and variability of the susceptibilities of the strong promoters is expected given that we tuned Λ^Topo^_*s*_ and Λ^Gyr^_*s*_ to capture these features. The correspondence nevertheless extends to the weakest promoters whose insensitivity to upstream context is reproduced without involving any additional fit. Furthermore, we also have the highest susceptibilities occurring for promoter strengths approximately three-fold lower than the maximum one. Even more significantly, although we constrained the unknown topoisomerase parameters based on the values of susceptibilities measured at a single upstream distance *d* = 3205 bp, our model quantitatively reproduces the full dependence of upstream susceptibility as a function of upstream distance. This is illustrated in Figure 5B where we show how we can find values of *k_b_, k_o_* and *σ_o_* for each of the three promoters studied in Figure 3 so as to reproduce the full dependence of their susceptibility as a function of upstream distance. These values are in fact tightly constrained (Fig. S8). For instance, good matches between experimental and theoretical results across all upstream distances as observed in Figure 5B impose to respectively use *σ_o_* = − 0.04 and *σ_o_* = − 0.05 for the weak and medium promoters (Fig. S8). This suggests that our approach may be used to infer promoter parameters.

**Figure 5:**
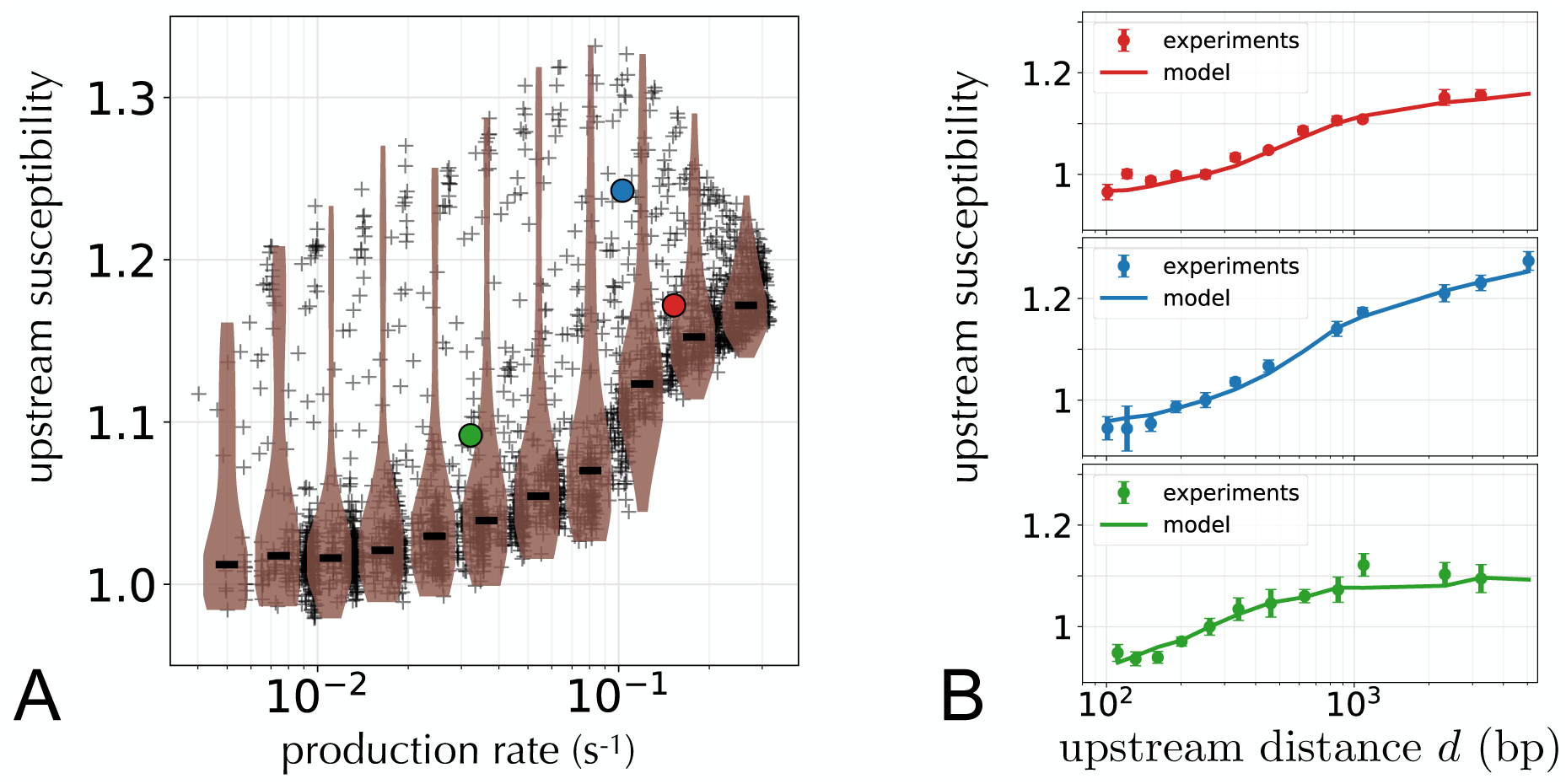
Upstream susceptibilities in the biophysical model – **A.** Susceptibility to upstream context versus promoter strength for the range of parameters indicated in Table 1. Horizontal lines of the violin plots indicate median values. **B.** Upstream susceptibility as a function of the upstream distance obtained in experiments (i.e. results of Fig. 3B) compared to the same quantity obtained in our model for three promoters indicated by colored dots in panel A (see Methods for the values of parameters).

#### 3.4.3 **Explaining promoter variability**

The dependence of transcript production on the upstream distance reflects antagonist effects that TopoI has on elongation and initiation. TopoI activity is indeed necessary to rescue RNAPs from stalling, and therefore enable elongation, but this activity causes the supercoiling density to jump by finite amounts, which can generate an “excess” of positive supercoiling that inhibits initiation by repressing OC formation (Fig. 6A). This inhibitory effect is prevalent at short upstream distances when TopoI activity induces strong variations of upstream DNA supercoiling density that cause the supercoiling density to be frequently above *σ_o_*, the threshold above which OC formation is prevented. In contrast, variations of TopoI-generated supercoils are dampened by a long upstream distance, with no impact on OC formation (Fig. 6B).

**Figure 6:**
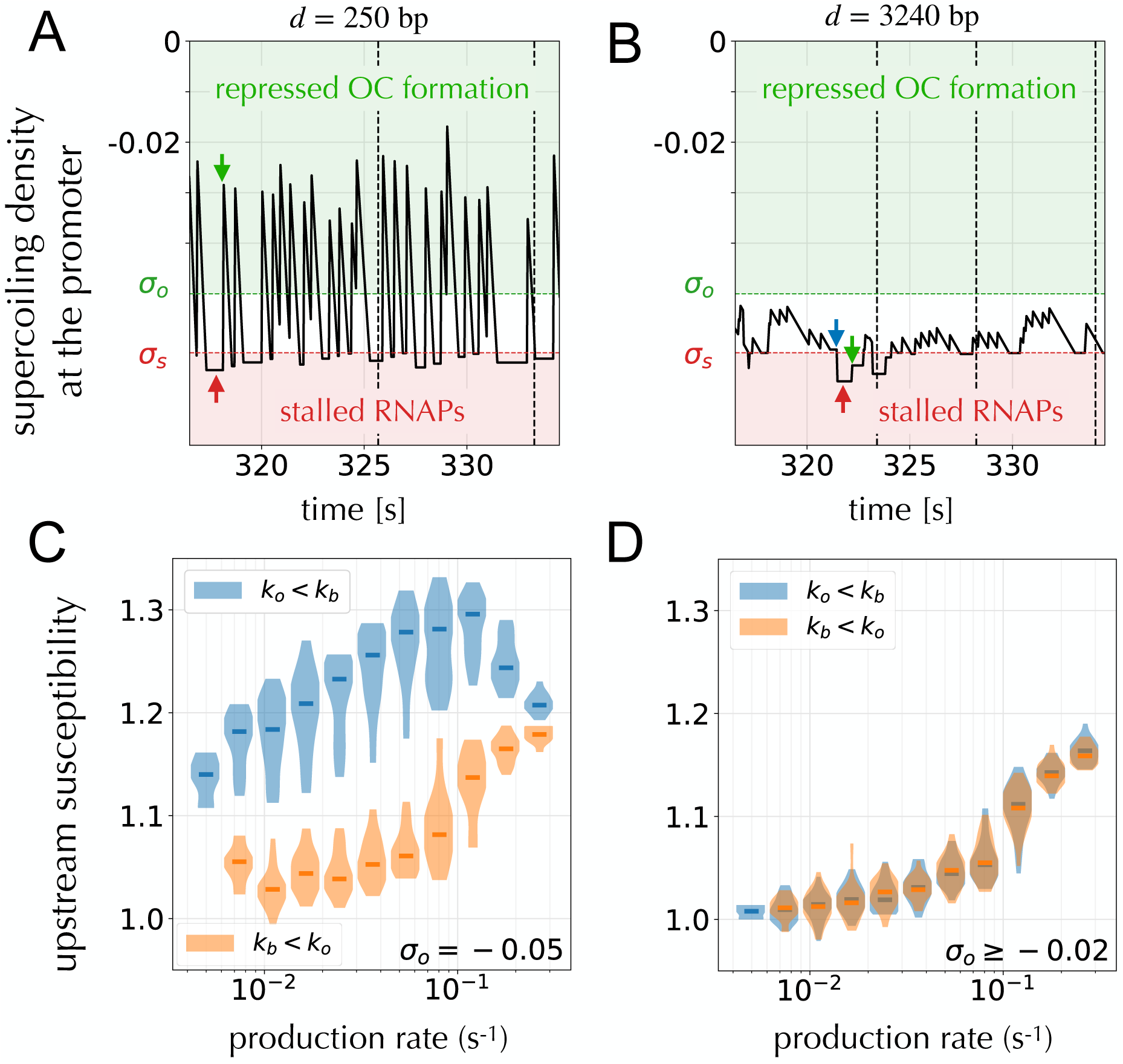
Simulation results rationalizing upstream susceptibilities – **A.** Values of the DNA supercoiling density at the promoter over a window of time in the stationary regime for short upstream distance (*d* = 250 bp) and a promoter corresponding to the medium promoter of Fig. 5B. The large positive jumps are the consequence of TopoI adding one supercoil (green arrow), while decreases are induced by RNAP translocations up to points where DNA supercoiling density is equal to the stalling threshold (*σ_s_*, red dashed line; the red arrow indicates RNAP stalling). When the upstream supercoiling density is above *σ_o_* (green dashed line), OC formation is repressed, preventing new initiations (vertical black dashed lines). **B.** Same as in panel A but for a long upstream distance (*d* 3200 bp), in which case positive supercoils added by TopoI are damped and the upstream supercoiling density remains below *σ_o_*. The blue arrow indicates a non-specific action of gyrase. **C.** Same as in Fig. 5A but considering promoters with *σ_o_* = 0.05 and distinguishing between those limited by binding (*k_b_ < k_o_*, in orange) and those limited by OC formation (*k_o_ < k_b_*, in blue). **D.** Same as in panel C but considering promoters with *σ_o_* 0.02, showing no difference between binding-limited promoters and OC formation-limited promoters.

We may also understand how promoters with comparable strength can respond differently to the presence of an upstream barrier by considering two underlying time scales: *k*^−1^, the time-scale of promoter binding, and *k*^−1^, the time-scale of OC formation. Indeed, while promoter strength depends roughly symmetrically on *k_b_* and *k_o_*, DNA supercoiling has a direct impact only on OC formation. Promoters with *k_b_ < k_o_*, i.e., limited by OC formation rather than by binding, are therefore expected to be more sensitive to changes of the upstream distance than those with *k_o_ < k_b_* (Fig. 6C). This effect depends on the value of *σ_o_*, as the lower *σ_o_*is, the more likely it is for the activity of TopoI to prevent OC formation. Here, in agreement with promoter supercoiling densities typically not exceeding 0.02 for an upstream distance *d* = 250 bp (Fig. 6A), differences in upstream susceptibility between promoters with limiting OC formation and those with limiting binding are manifest only when *σ_o_ < −*0.02 (Fig. 6D).

## 4 Discussion

Gene context is recognized as an important determinant of gene expression with several possible mechanisms at play, including local concentration effects, transcriptional read-through, RNAP interferences and DNA supercoiling. It is generally unknown, however, which mechanism – if any – is prevalent in given *in vivo* conditions. A major impediment has been the absence of data from *in vivo* experiments where the gene context is fully controlled. Here we introduced an insulated genetic system that realizes *in vivo* the simplest case, also known as the twin transcriptional-loop model: a single gene transcribed on a DNA segment delimited by two topological barriers. The study of this minimal system suggests that DNA supercoiling is a prevalent mechanism via which genetic contexts affects expression *in vivo*. It also allows us to assess how DNA supercoiling is handled *in vivo* and how it affects gene expression. We find expression rates to be limited by the presence of an upstream topological barrier but not of a downstream topological barrier. The larger the distance to the upstream distance, the larger the expression rate but the susceptibility of a gene depends non-linearly on the distance and is strongly promoter dependent.

To interpret our experimental results, we developed a first-principle biophysical model of transcription with no free parameter but the mode of action of TopoI and gyrase and the values of promoter parameters. In this model, RNAP elongation generates DNA supercoiling on each side of the elongating RNAP, which in turn affects the elongation of other RNAPs as well as OC formation during initiation. Specifically, downstream positive supercoiling inhibits elongation while upstream negative supercoiling inhibits OC formation. DNA supercoiling on both ends of the gene is then modulated by the action of TopoI and gyrase which we considered to be either non-specific, i.e., scaling with the size of the domain, or specific, i.e., localized at the start or end of the gene. We find the model to account for our experimental data only when TopoI and gyrase are allowed to act specifically. In line with previous works on gyrase [32, 60] and TopoI [60, 61, 36] in various bacteria, including *E. coli*, our results thus demonstrate that these topoisomerases are essential facilitators of transcription. Our analysis further reveals that the removal rate of supercoils is lower for TopoI than for gyrase. We also find that elongating RNAPs produce supercoils at a rate higher than that of TopoI removing negative supercoils. Altogether, our findings therefore show that elongation is mainly controlled by TopoI activity.

While TopoI enables elongation, simulations of our biophysical model reveal an additional antagonistic effect at the core of the large upstream susceptibilities: TopoI represses initiation when the distance to the upstream barrier is too short to dampen changes in supercoiling density. This antagonism is topologically inevitable due to the discrete nature of the supercoils that TopoI adds, which unavoidably translate into discrete increases of the supercoiling density whose size is all the larger that the upstream distance to a topological barrier is short. Consistent with the predictions that TopoI plays a primary role in these phenomena and that the mechanisms involve localized variations of supercoiling at the promoter, inhibiting gyrase has minimal effect on upstream susceptibility, despite significant changes in average supercoiling density and gene expression (Figs. S17-S19). We also verify that removing or inactivating topoisomerases TopoIII and TopoIV, which are not accounted for in our model, has no impact on upstream susceptibility (Fig. S18).

Nevertheless, variability in upstream susceptibility is observed among strong promoters which reflects two distinct contributions to promoter strength that are differentially affected by TopoI-induced supercoiling: promoters limited by binding are nearly insensitive to upstream context, while those limited by OC formation are sensitive. Our model indicates that the latter occurs when both the OC formation rate *k_o_* is smaller than the binding rate *k_b_* and when *σ_o_*, the threshold over which OC formation is permitted, is sufficiently close to the RNAP stalling density, *σ_s_*. Predicting the susceptibility of a specific promoter therefore requires the three parameters *k_b_*, *k_o_* and *σ_o_* – to which in practice *k_e_* must be added, which is encompassed in *k_b_* in our model. While a systematic inference of these parameters is beyond the scope of the present work, our study of multiple promoters over a range of parameter values is already highly informative and constrains not only qualitatively but also partly quantitatively the activity of topoisomerases. We thus obtained an upper bound on the non-specific activity rate of TopoI, namely *λ*^Topo^_*ns*_ *<* 5.10^−4^ Lk.bp^−1^.s^−1^, as well as an expected range of values for the specific activity of both gyrA and TopoI, namely Λ^Gyr^_*s*_ 2 Lk.bp^−1^.s^−1^ and Λ^Topo^_*s*_ 1 2 Lk.bp^−1^.s^−1^, respectively. Moreover, reproducing quantitatively the full dependence of the sensitivity of specific promoters on upstream distances as in Fig. 5B strongly constrains the possible values of *k_b_*, *k_o_* and *σ_o_* and hence, provide, a promising road to estimate promoter parameters.

In bacteria as in eukaryotes, the dynamics of transcription of many genes is bursty, with phases of activity separated by long phases of inactivity [64]. This manifests as a non-Poissonian distribution of transcripts in cell populations, with a high proportion of cells containing very low numbers of transcripts [64, 65, 32]. *In vivo* experiments showed that these properties depend on gyrase activity [32]. In particular, gyrase under-expression (over-expression) leads to a higher (lower) proportion of cells with very few transcripts. This is in agreement with longer inactivity periods where DNA gyrase is absent and, hence, during which accumulated positive supercoiling blocks elongation [32]. The time scales between active and inactive phases is typically of the order of ten minutes [64, 65], much larger than those associated with the specific activity of the topoisomerases obtained in our work (on the order of a second). In its current form, our model does not account for these effects and a precise study to refine it is beyond the scope of this work. Nevertheless, we verify that adding a “slow” two-state (bound gyrase/unbound gyrase) mechanism to the here-obtained “fast” gyrase activity does not change qualitatively our findings, while capturing bursting properties similar to those observed *in vivo*, including the impact of gyrase concentration (Fig. S20). This illustrates how our results can both arise from a different mechanism than transcriptional bursting and be perfectly consistent with its occurence.

In future work, it will be interesting to further elaborate and test our model predictions by repeating our measurements with topoisomerases exhibiting different levels of activities, more particularly with TopoI. While this may be achieved through their under or over-expression, the use of mutants or of inhibitors, a fundamental difficulty should be noted: modulating the activity of one topoisomerase is expected to impact various cell parameters, including for instance the activity of other topoisomerases [16]. More generally, the balance between gyrase and TopoI activities determines the levels of supercoiling, nucleoid compaction, and viability in bacteria [66, 67, 68, 69]. As these global physiological changes are poorly understood from a quantitative perspective, relating the results of such experiments to those of our model predictions may be a non-trivial challenge.

From a genomic perspective, our system purposely defines a limit case where a single gene is fully insulated from other genes. In genomes, no gene is totally insulated from its neighbors but different nucleoid-associated proteins as well as RNAPs themselves may isolate larger groups of genes. In future work, our system could be scaled up to insulate two and more genes and therefore provide valuable information on the consequences of genome organization for gene regulation. In any case, studying the feedback of a single transcribed gene onto itself is a pre-requisite to studies with more intricate gene contexts, as well as a proof-of-concept of their interest. In particular, our findings underscore the need to model topoisomerase activity accurately in order to achieve a quantitative understanding and prediction of the behavior of gene expression, whether individual or collective.

Additionally, our results are of interest for synthetic biology as they demonstrate a mechanism by which gene expression can be finely controlled. The modulation of gene expression by the distance to an upstream is indeed robust, i.e., independent on the composition of the sequence separating the gene to the topological barriers (Fig. S9), and its simple monotonous dependence is remarkable when contrasted with the complex dependence to promoter sequences (Fig. S2). This is all the more remarkable that the effects are comparable in magnitude to modifying the up-element sub-structure of a promoter (Fig. S10).

Identifying which effects are robust and therefore amenable to an explanation and to experimental control is essential both to the theory and the engineering of biological processes. Counterintuitively, our results suggest that, for transcription, gene context may be more amenable to quantitative explanations and experimental control than promoter sequences despite involving long-range indirect coupling between DNA and RNA polymerases.

## Data and code availability

The experimental data and a Python code for reproducing the figures that represent them is available at https://zenodo.org/record/8174873 and a Python implementation of the biophysical model is available at https://zenodo.org/record/8167497.

## Acknowledgments

We thank Estelle Crozat, Hans Geiselmann, Călin Guet, Thomas Hindré and Bianca Sclavi for fruitful discussions. We acknowledge funding from FRM AJE20160635870, IRIS OCAV Idex ANR-10-IDEX-0001-02 PSL, ANR-18-CE12-0012, LabEx MemoLife and PSL-Qlife.

