## Supplementary information for "Assessing *in vivo* the impact of gene context on transcription through DNA supercoiling"

#### Contents

**1 Supplementary text** **1**
1.1 Inference of gene expression rate from microplate experiments . . . 1
1.2 Processing of microplate reader data . . . 2

**2 Supplementary Tables** **4**

**3 Supplementary Figures** **5**

### 1 Supplementary text

#### 1.1 Inference of gene expression rate from microplate experiments

The rate at which a test gene is transcribed depends not only on its promoter and genetic context, but also on the physiological state of the cell, which includes for instance the number of available ribosomes [1]. We are interested in comparing transcription rates between systems with varying promoters, genetic constructs and conditions (e.g., in presence or not of IPTG). All these factors can affect the physiology of bacteria and, therefore, the estimation of transcription rates. To compare intrinsic transcription rates where extrinsic physiological effects are factored out, the common approach is to normalize the transcription of the test gene by the transcription of a control gene [2]. This corresponds to taking as a measure of transcription rate

$$\alpha_{\text{test}} = \frac{dF_t^{\text{test}}/dt}{dF_t^{\text{control}}/dt} \tag{S1}$$

where $F_t^g$ denotes the fluorescence of proteins produced by gene $g$ at time $t$. In addition to assuming fast-folding and slow-degrading fluorescent proteins, this approach rests on three hypotheses:

(H1) Changes in physiology impact transcription multiplicatively so that $dF_t^{\text{test}}/dt = a_{\text{test}}\Phi$ and $dF_t^{\text{control}}/dt = a_{\text{control}}\Phi$ with $a_{\text{test}}$ and $a_{\text{control}}$ independent of a global physiological variable $\Phi$ (defined here at the population level and therefore depending on the population size $N$). This guarantees that $\alpha_{\text{test}} = a_{\text{test}}/a_{\text{control}}$ does not depend on $\Phi$.

(H2) An interval of times can be found over which the ratio $\alpha_{\text{test}}$ is approximatively constant.

(H3) The transcription of the control gene is independent of that of the test gene so that $a_{\text{control}}$ is independent of $a_{\text{test}}$. This guarantees that changes in $\alpha_{\text{test}}$ are proportional to changes in $a_{\text{test}}$. On the other hand, $\Phi$ may depend on $a_{\text{test}}$ and $a_{\text{control}}$.

Hypothesis (H1) is the fundamental assumption that intrinsic and extrinsic (physiological) factors can be separated. Hypothesis (H2) is expected to be verified in the exponential phase of bacterial growth where the instantaneous growth rate $(dN/dt)/N$ is constant. In this case, we expect constant transcription rates per bacterium $(dF_t^{\text{test}}/dt)/N$ and $(dF_t^{\text{control}}/dt)/N$ and therefore a constant ratio $\alpha_{\text{test}}$. Hypothesis (H3) is expected to be verified when the genes are constitutive and belong to distinct genetic contexts.

In our experiments as in other experiments where measurements are made in a microplate reader [3, 4], no exponential phase is observed (Fig. S11). Besides, to demonstrate that our genetic design with a closed topological loop (Fig. 1B) insulates the test (upstream) gene from the control (downstream) gene, we cannot rely on hypothesis (H3) which assumes that this is the case. Instead, we need a reproducible measure of transcription involving only the gene of interest (test or control) which can be derived from dynamical measurements of gene fluorescence and optical density with cells that are not growing exponentially.

This problem has been faced in previous studies where two approaches have been applied, based on the quantity $(dF^{\text{test}}/dt)/N$. In one approach, this quantity was taken at its maximum [3], while in the other is was taken at the maximum of instantaneous growth rate $(dN/dt)/N$ [4]. These approaches capture biologically relevant differences in gene expression but have two shortcomings. Fundamentally, they are not easily justified as they do not refer to any underlying stationary process (Fig. S12A). Practically, we found them to be insufficiently precise for our purpose, where differences in transcription rate of the order of 10% are of interest. To derive an alternative measure, we reasoned that bacterial growth is itself dependent on the physiological state and may therefore serve as a control quantity. Assuming $dN/dt = \gamma\Phi$ with $\gamma$ independent of $\Phi$ and $a_{\text{test}}$, this suggests to consider

$$\beta_{\text{test}} = \frac{dF_t^{\text{test}}/dt}{dN/dt}. \tag{S2}$$

Consistent with our hypotheses, we observe that this quantity is contant over an interval of times preceding the decrease of the instantaneous growth rate $(dN/dt)/N$ (Fig. S12C). Further, we verify that the quantity $\alpha'_{\text{test}}$ defined over this interval is essentially proportional to the quantities defined in previous approaches (Fig. S14). It is, however, more precise, i.e., more reproducible (Fig. S15).

In Fig. 1D of the main text, where we compare open and closed contexts, we therefore take $\beta_{\text{test}}$ and $\beta_{\text{control}}$ from Eq. (S2) to report the transcription rate of either the test or control gene. The other approaches, while less precise, lead to results that are perfectly consistent with our conclusion, namely that closing the topological loop defines an insulated gene context (Fig. S16). In Figs. 2, 3 and 4 of the main text we rely on this result which justifies hypothesis (H3) and report $\alpha_{\text{test}}$ from Eq. (S1) averaged over the interval of time where this quantity is approximatively constant (Fig. S12A).

#### 1.2 Processing of microplate reader data

The raw temporal data for the optical density and fluorescence is linearly interpolated over 750 points, from the $\sim 50$ raw data points using the interp1d module from the SciPy library in Python. The interpolated data is then filtered using a 2$^{\text{nd}}$ order polynomial by a Savitzki-Golay filter using the savgol module from the SciPy library in Python using window size of 101. The codes are available as Jupyter notebooks provided as Supplementary Information.

The relative differences in gene expression rates are only weakly sensitive to the exact parameters used for interpolation and filtering. However, the use of different parameters can lead to quantitative differences when different growth environments are used. We systematically applied to all datasets the same set of interpolation and filtering parameters as well as the time duration over which the signal is temporally averaged. These parameters were chosen so as to minimise the differences in inferred expression rates between replica from a given genetic construct. In order to focus the analysis on the bulk of the signal as opposed to the very beginning (dominated by measurement noise) or the end (saturation phase), the data analysis code also makes use of a parameter controlling how far from the time of maximum growth rate can the data be considered. The time window of stable signal (over which the temporal averaging is performed) is independently determined for $(dF_t^{\text{upstream}}/dt)/(dN_t/dt)$, $(dF_t^{\text{downstream}}/dt)/(dN_t/dt)$ and $(dF_t^{\text{upstream}}/dt)/(dF_t^{\text{downstream}}/dt)$, so as to not impose any assumption upon protein folding time.

#### 2 Supplementary Tables

| Label | Name | Sequence (aligned by -10 element) |
|---|---|---|
| p01 | apFAB67 | TTGACATCAGGAAAATTTTTCTG**CATAAT**TATTTCATATCAC |
| p02 | pR | TATCTAACACCGTGCGTGTTGACTATTTTACCTCTGGCGGT**GATAAT**GGTTGCA |
| p03 | pTet | TCCCTATCAGTGATAGAGATTGACATCCCTATCAGTGATAGA**GATACT**GAGCACATCAGCAGGACGCACTGACC |
| p04 | apFAB40 | AAAAAGAGTATTGACTTCAGGAAAATTTTTCTG**TATAAT**GTGTGGATGTTCA |
| p05 | apFAB45 | AAAAAGAGTATTGACTTCGCATCTTTTTGTACC**TATAAT**GTGTGGATAGCGG |
| p06 | apFAB93 | AAAAAATTTATTTGCTTTCAGGAAAATTTTTCTGTAG**ATTTAA**CGTATGCCCA |
| p07 | apFAB101 | AAAAAATTTATTTGCTTTTTATCCCTTGCGGCGA**TATAAT**AGATTCATCTTAG |
| p08 | apFAB341 | TTGACAATTAATCATCCGGCTCG**TAATAT**GTGTGGATGGTT |
| p09 | apFAB79 | AAAAAATTTATTTGCTTTAAAGTCTAACCTATAG**GATACT**TACAGCCATAGTCT |
| p10 | BBa J23119 | TTGACAGCTAGCTCAGTCCTAGG**TATAAT**GCTAGCACGAA |
| p11 | apFAB61 | TTGACAATTAATCATCCGGCTCG**TATAAT**AGATTCATTAGAG |
| p12 | apFAB70 | TTGACATCGCATCTTTTTGTACC**TATAAT**GTGTGGATAGAGT |
| p13 | parcB | CAACGGAGTAGGTCGTTGAGGGGAATTCCGCATTTCTCACACAATTTATAACGTAACTGTCAGAATTGGG**TATTAT**TGGGGC |
| p14 | pkdsA | GCTGGACGTATGGTTAAAGGGAAATATCAGCCCGTCGGCGGAACTGTTTTATGAAGACCTTGATGAAGCT**GATAAC**ATTGAG |
| p15 | prna | TACGTTCGCCAAACGCTGAATAAATCATCACGCCCGCCAGGTAAGCCCACCTGGCGGGCTTTTTTATGA**TTTAAT**AGATAGT |
| p16 | pyacL | AATGAGATTCCCGGGCATTTTTTTATTTCTAAACCATCGCCGTTCCGCTGTTTTTCTCCGGTAAGGCTGC**GATAAT**TACATC |
| p17 | apFAB73 | TTGACATCGCATCTTTTTGTACCTAG**ATTTAA**CGTATTCGGC |
| p18 | J23101 | TTTACAGCTAGCTCAGTCCTAGG**TATTAT**GCTAGCCAGTT |
| p19 | J23100 | TTGACGGCTAGCTCAGTCCTAGG**TACAGT**GCTAGCTTAAT |
| p20 | apFAB47 | TTGACAATTAATCATCCGGCTCT**TAATAT**GTGTGGACGAGG |
| p21 | apFAB66 | TTGACATCAGGAAAATTTTTCTG**TATAAT**AGATTCATCTCAA |
| p22 | apFAB77 | TTGACATTTATCCCTTGCGGCGA**CATAAT**TATTTCATTTTGG |
| p23 | apFAB67* | TTGACATCAGGAAAATTTTTCTG**CATAAT**CCCGGCATATCAC |
| p24 | apFAB67** | TTGACATCAGGAAAATTTTTCTG**CATAAT**GTGTGGATATCAC |
| p25 | apFAB67*** | TTGACATCAGGAAAATTTTTCTG**TAATAT**TATTTCATATCAC |
| p26 | apFAB341* | TTGACAATTAATCATCCGGCTCG**TAATAT**TATTTCATGGTT |
| p27 | apFAB341** | TTGACAATTAATCATCCGGCTCG**TAATAT**CCCGGCATGGTT |
| p28 | apFAB341*** | TTGACAATTAATCATCCGGCTCG**CATAAT**GTGTGGATGGTT |
| | lacO barrier | GCTTATGACGACAAGAATTGTGAGCGGATAACAATTGCACAGAATACATTATGAAGTCTCGAATTGTGAGCGGATAACAATTGCCCTGAGTACAGCTG |

Table S1: List of promoters used in our experiments. Fig. 1D reports data with the 10 promoters labeled from p02 to p12, Fig. 2A with the 19 promoters labeled from p01 to p20 with the exception of p14, Fig. 2B with the 28 promoters labeled from p01 to p28, Fig.3 with the 3 promoters p01 (in blue), p05 (in green) and p12 (in red), Fig. S2B-D with the 2 promoters p01, p08 and their 6 variants p23 to p27.

| | |
|---|---|
| Terminator B0014 | TCACACTGGCTCACCTTCGGGTGGGCCTTTCTGCGTTTATATACTAGAGAGAGAA TATAAAAAGCCAGATTATTAATCCGGCTTTTTTATTATTT |
| Terminator T1 | GGCATCAAATAAAACGAAAGGCTCAGTCGAAAGACTGGGCCTTTCGTTTTATCTG TTGTTTGTCGGTGAACGCTCTCCTGAGTAGGACAAATCCGCCGCCCTAGA |
| mCerulean RBS (apFAB837) | ATCTTAATCTAGCAGGGGATATTTT |
| mVenus RBS | GGTTGCATGTACTAGAGTTCATTAAAGAGGAGAAAGGTACC |

Table S2: Sequences of the two terminators and two ribosome binding sites (RBS) used in our constructs (see Methods).

### 3 Supplementary Figures

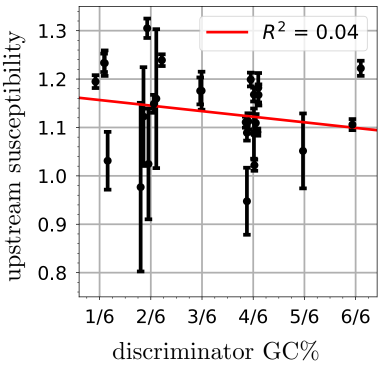

Figure S1: Upstream susceptibility versus GC content of the discriminator sequence for the promoters of Fig. 2B (a small random component is added to the value of GC content to limit the overlap between data points). A linear regression (red line) results in a coefficient of determination $R^2 = 0.04$ with a p-value $p = 0.3$ (two-sided p-value for a hypothesis test whose null hypothesis is that the slope is zero using Wald Test with t-distribution of the test statistic), indicating no significant correlation.

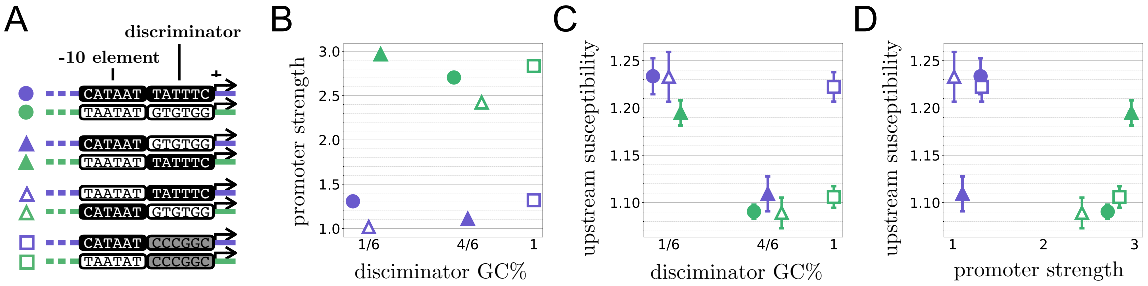

Figure S2: Dependence of upstream susceptibility on the GC content of the 6 nucleotides preceding the start codon, which overlaps with the discriminator. – **A.** We took two promoters (purples and green full circles) and swapped either these 6-nucleotide sequences (full triangles) or their -10 elements (empty triangles). We also made variants where the GC content of the 6-nucleotide sequence is maximized (empty squares). **B.** Promoter strength of each construct versus the GC content of the 6-nucleotide sequence. **C.** Upstream susceptibility of each construct versus the GC content of the 6-nucleotide sequence. The original purple promoter is significantly more susceptible than the original green promoter (full circles). Swapping their 6-nucleotide sequence effectively swaps their susceptibilities (full triangles), while swapping their -10 elements has no effect (empty triangles). Maximizing the GC content of their 6-nucleotide sequence has, however, little effect on their susceptibility (empty squares), showing that susceptibility of a promoter to upstream context is not simply related to the GC content of its discriminator. **D.** Upstream susceptibility of each construct versus promoter strength.

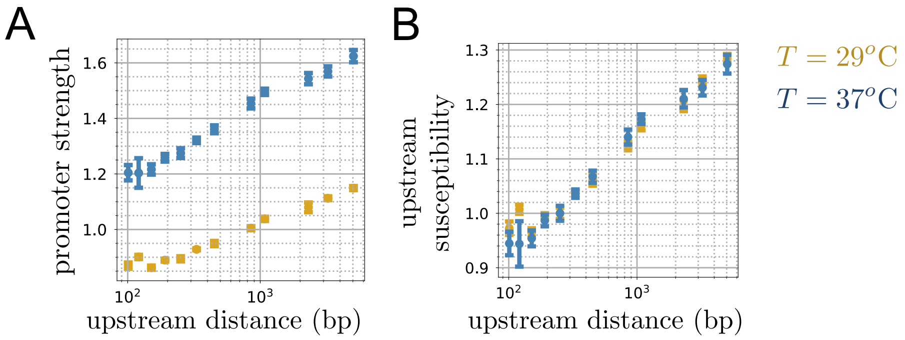

Figure S3: Dependence on temperature of promoter strength and upstream susceptibility for the intermediate (blue) promoter of Fig. 3. The data in Fig. 3 was collected at $T = 37^o$C. It is compared here to data collected at $T = 29^o$C (in yellow), showing that promoter strength is temperature dependent but not upstream susceptibility.

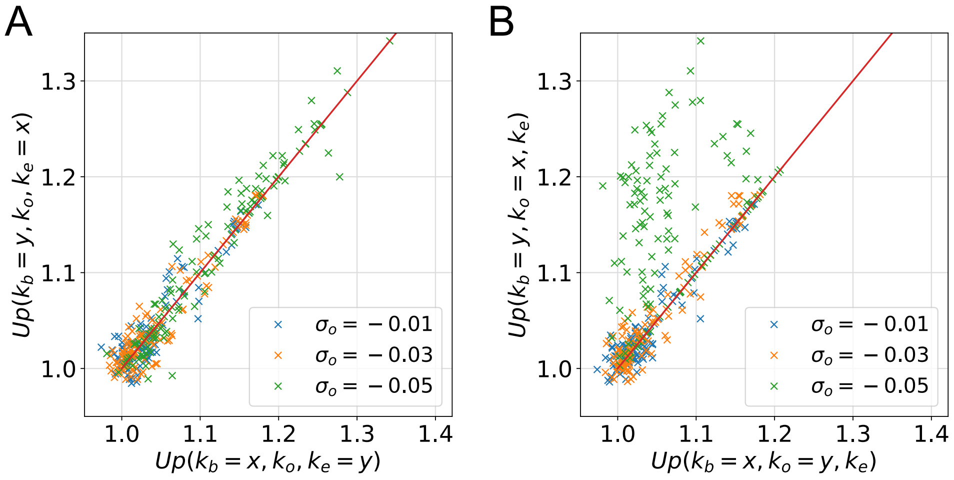

Figure S4: Redundant impact of $k_b$ and $k_e$ on upstream susceptibilities. **A.** Correspondence between upstream susceptibilities when exchanging the values of $k_b$ and $k_e$, showing a symmetric effect of $k_b$ and $k_e$. **B.** Correspondence between upstream susceptibilities when exchanging the values of $k_b$ and $k_o$, showing, in contrast, a non-symmetric effect of $k_b$ and $k_o$. To generate these plots, we considered 648 combinations of values of $k_b$, $k_o$, $k_e$ and $\sigma_o$ where the frequencies were taken in $\{0.01 \times 4^i\}_{i=0,..,5}$ and $\sigma_o$ in $\{-0.05, -0.03, -0.01\}$.

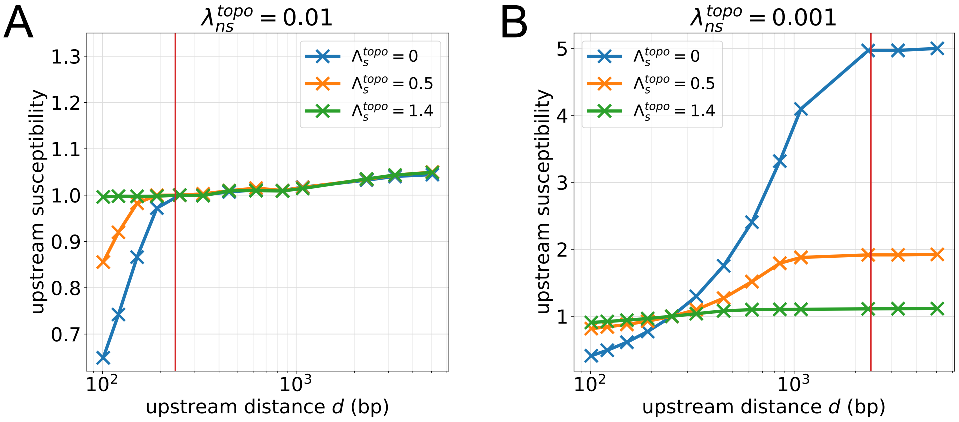

Figure S5: Effect of $\lambda_{ns}^{\mathrm{Topo}}$ on upstream susceptibility. **A.** Susceptibility to upstream context versus upstream distance for a situation where $\lambda_{ns}^{\mathrm{Topo}} = 0.01$ Lk.bp$^{-1}$.s$^{-1}$ for three different values of $\Lambda_s^{\mathrm{Topo}}$ and for $\Lambda_s^{\mathrm{Gyr}} = -2.5$ Lk.s$^{-1}$, showing a first increasing regime up to a characteristic length on the order of $v_m/(n\lambda_{ns}^{\mathrm{Topo}})$ (vertical red line). **B.** Same as panel A but with $\lambda_{ns}^{\mathrm{Topo}} = 0.001$ Lk.bp$^{-1}$.s$^{-1}$.

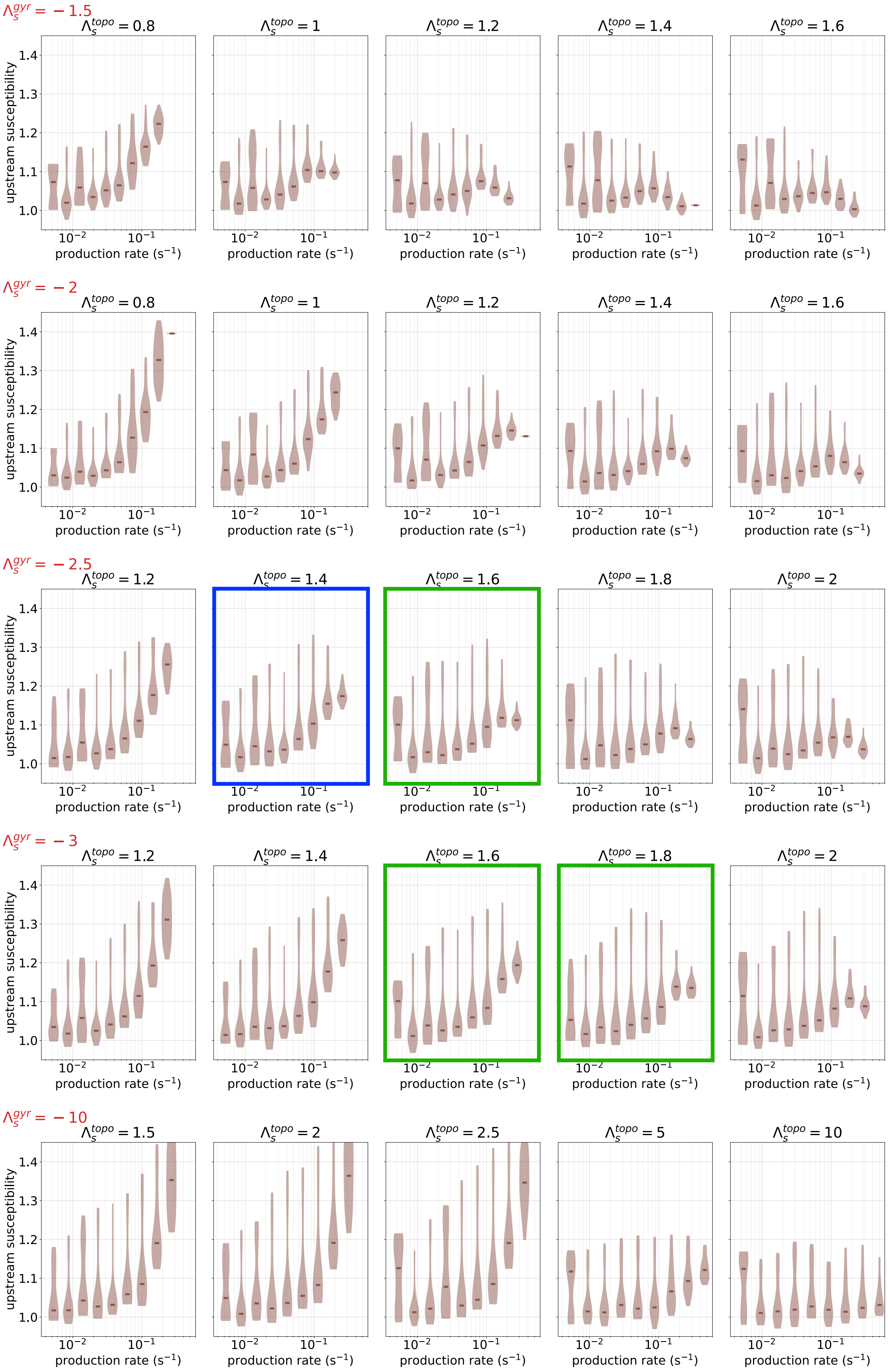

Figure S6: Susceptibilities to upstream context of the range of promoters indicated in Table 1 when considering different values of $\Lambda_s^{\mathrm{Topo}}$ and $\Lambda_s^{\mathrm{Gyr}}$. From top to down, the rows correspond to $\Lambda_s^{\mathrm{Gyr}} = -1.5, -2, -2.5, -3, -10$ Lk.s$^{-1}$. The panel highlighted in blue corresponds to the combination of parameters used in the main text (Fig. 5A). Panels highlighted in green are additional combinations compatible with our experimental results, considering a maximum susceptibility at $\sim 1.3$ and the susceptibility of the strongest promoters typically lying between 1.1 and 1.2.

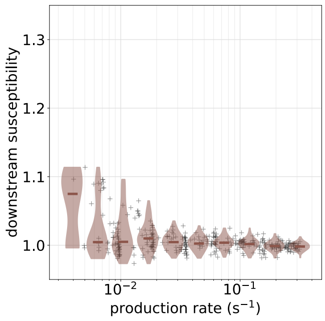

Figure S7: Susceptibility to downstream context versus promoter strength for the range of parameters indicated in Table 1 of the main text. Horizontal lines of the violin plots indicate median values. The range of the y-axis is the same as in Fig. 5A of the main text.

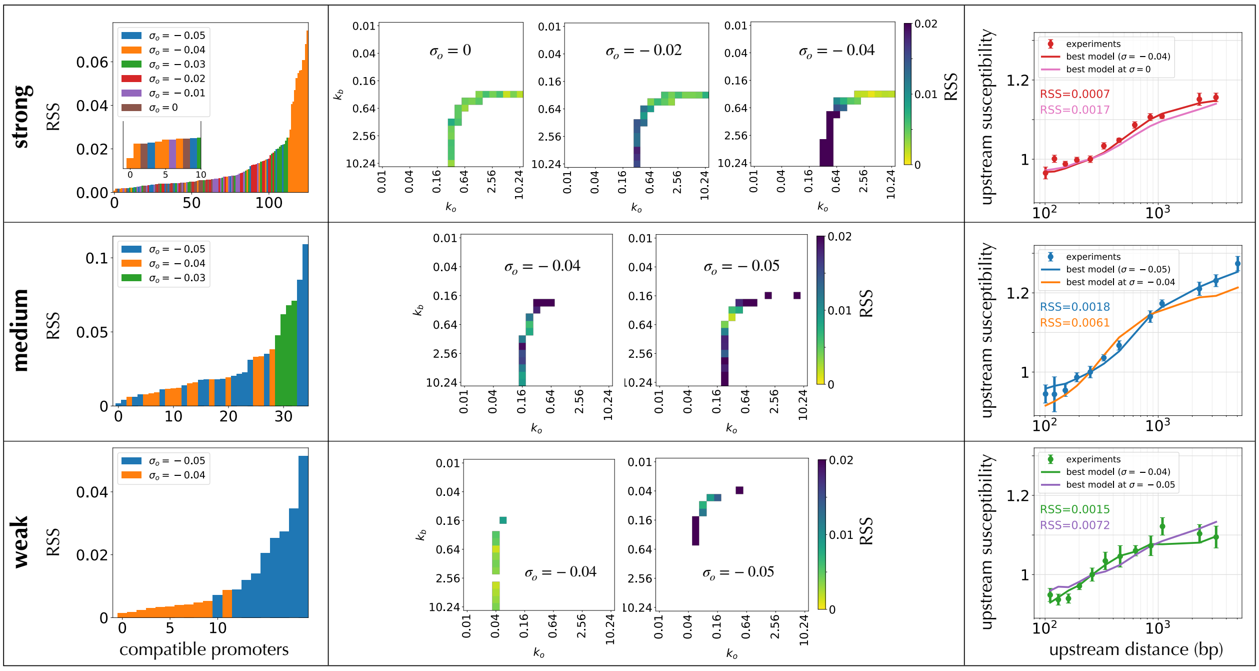

Figure S8: Inference of best parameters for fitting the dependence of the upstream susceptibility as a function of the upstream distance in Figure 5B – We first consider all promoters with a production rate in the intervals $0.033\,\mathrm{s}^{-1} \pm 10\%$ (weak promoters), $0.1\,\mathrm{s}^{-1} \pm 10\%$ (medium promoters) and $0.16\,\mathrm{s}^{-1} \pm 10\%$ (strong promoters). Production rates of medium promoters are typical of those of maximal upstream susceptibility (see Fig. 5A in the main text), while the two other intervals are chosen to reproduce the ratio of production rates among the strong, medium and weak promoters as observed in the experiments (see Fig. 3A in the main text). We quantify the goodness of fit with the residual sum of squares (RSS) when comparing the data to model predictions for the strong (top row), medium (middle row) and weak (bottom row). The leftmost panels show promoters' RSS by sorting the promoters according to this value. The bars are painted according to the value of the promoter's $\sigma_o$; this reveals strong constraints on the value of $\sigma_o$ for the weak and medium promoters. The middle panels show the RSS for a given $\sigma_o$ as a function of the two other promoter parameters, $k_b$ (y-axis) and $k_o$ (x-axis); the parameters within a one-dimensional manifold corresponding to $1/k_b + 1/k_o$, the promoter strength. The rightmost panels report the dependence of the upstream susceptibility as a function of the upstream distance for the best model as well as for the best model with the second best value of $\sigma_o$.

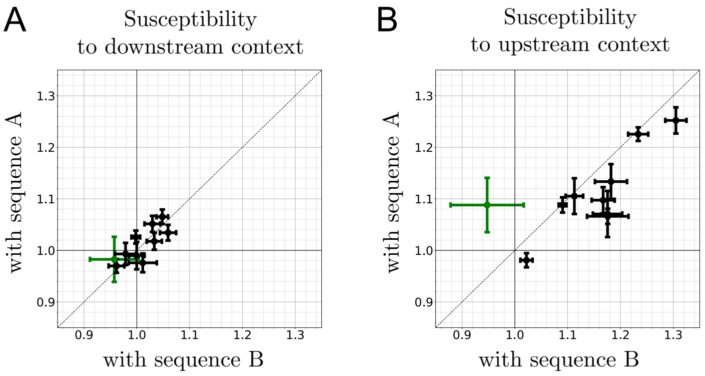

Figure S9: Replicate of measurements in Fig. 2 with different downstream and upstream sequences – **A.** When replacing the 3 kb downstream sequence (sequence A) by another sequence (sequence B), we verify that the downstream susceptibility remains below 10%. The effects on the order of 5% observed for some promoters appear, however, to be reproducible. Sequences A and B are respectively labeled uL and dC in the file Distances.csv provided as Supplementary Data. **B.** When replacing the 3 kb upstream sequence (sequence A) by another sequence (sequence B), we verify that the upstream susceptibilities are strongly correlated, indicated that the upstream susceptibility is largely sequence independent. The main outlier (represented in green) may be explained by a difference of up-element (see Fig. S10).

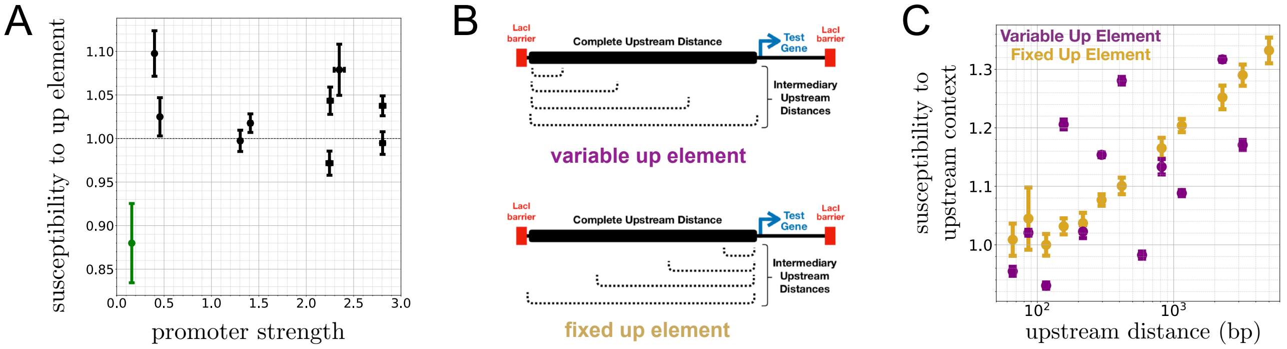

Figure S10: Contribution of the up element to upstream susceptibility – **A.** A susceptibility to the up element is obtained here as the ratio of expression rate for two constructs that differ only by their up element – up element of sequence A over up element of sequence B with in both cases short upstream and downstream distances. The main outlier in Fig. S9 is also the one with largest susceptibility to the up element (in green), indicated that the up-element is the likely origin of the difference. **B.** The data of Fig. 3 corresponds to upstream sequences that differ in length but have the same up element (design indicated as "fixed up element"). For comparison, we also acquired data with reversed sequences, which have different up elements (design indicated as "variable up element"). **C.** Varying the sequence up element is found to produce effects comparable to varying the sequence length. This data was acquired at 29° C and the data for a Fixed Up Element therefore corresponds to the data shown in Fig. S3 at this temperature.

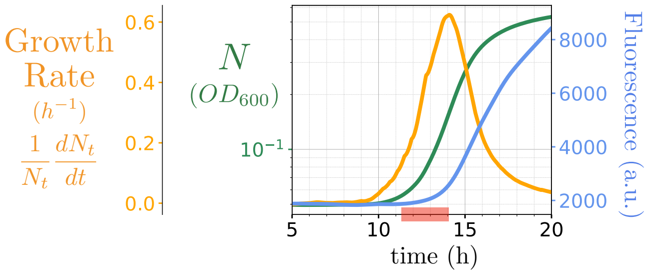

Figure S11: Typical time series from microplate reader experiments – Shown are the fluorescence from the test gene (in blue), the optical density ($OD_{600}$) reporting the cell density $N$ (in green) and the instantaneous growth rate $(dN/dt)/N$ computed from the derivative of the OD (in yellow). This instantaneous growth rate is never constant over the dynamic range of the experiment, indicating an absence of exponential growth phase. The promoter used here and in Fig. S12 is apFAB45 in Table S1 (also labeled p05_uNdB_i0_t37 in the datasets). The red segment on the x-axis indicates the time-window over which we estimate the transcription rate for this particular experiment, as explained in Fig. 12A.

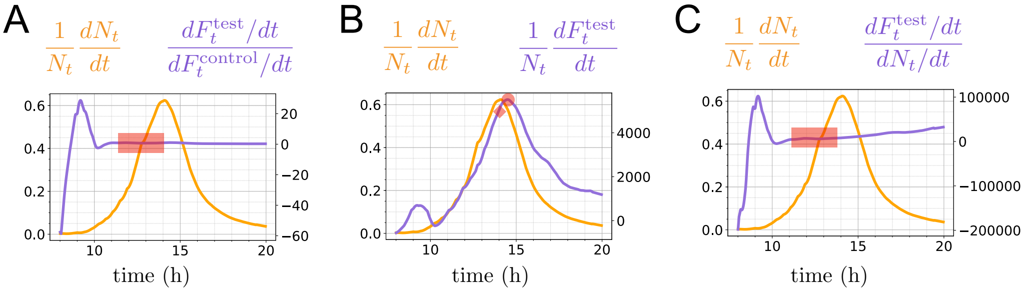

Figure S12: Illustration of different approaches to measure transcription rate from the data shown in Fig S11 – **A.** When a control gene is present, the ratio $(dF_{\text{test}}/dt)/(dF_{\text{control}}/dt)$ (in purple) is approximatively constant over an extended time interval. Averaging this value over a time window prior to the entrance into stationary phase (in red) provides a measure of transcription rate that we use in all figures of the main text, except Fig. 1D. **B.** In absence of a control gene, it has been proposed to consider $(dF_{\text{test}}/dt)/N$ which corresponds for exponentially growing cells to a rate per cell but which is here never constant, and to retain either its value at its maximum [3] (red circle) or at the point of maximal instantaneous growth [4] (red diamond). **C.** An alternative that we propose is to consider $(dF/dt)/(dN/dt)$ where $(dF/dt)/N$ is normalized by $(dN/dt)/N$. This quantity is nearly constant over a time window preceding the entry in stationary phase. Averaging over this window results in a reproducible measure that does not require a control gene (Fig. S15). This is the approach that we take in Fig. 1D.

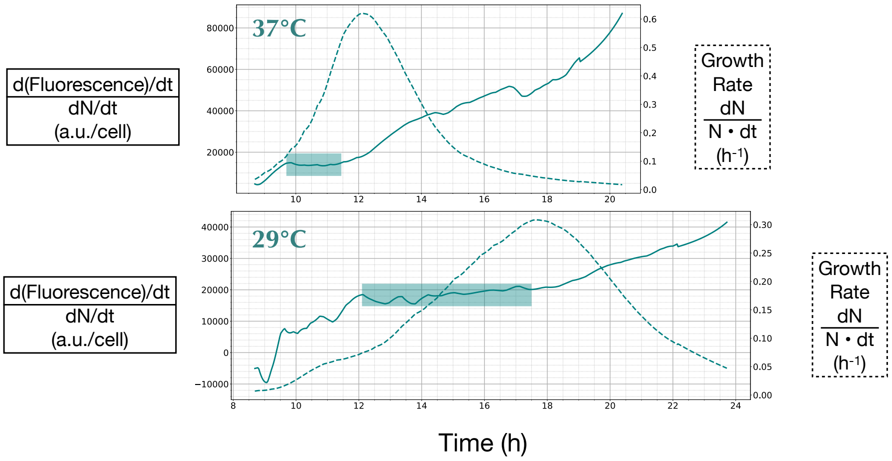

Figure S13: Lowering the temperature from 37°C to 29°C slows down population growth (dotted line representing $(dN_t/dt)/N_t$) and extends the time window over which the ratio $(dF_t^{\text{test}}/dt)/(dN_t/dt)$ (full line) is nearly constant from under 2h to over 5h. These curves were obtained for one clone with promoter p01 (Table S1), upstream distance uC (116 bp) and downstream distance dB (507 bp).

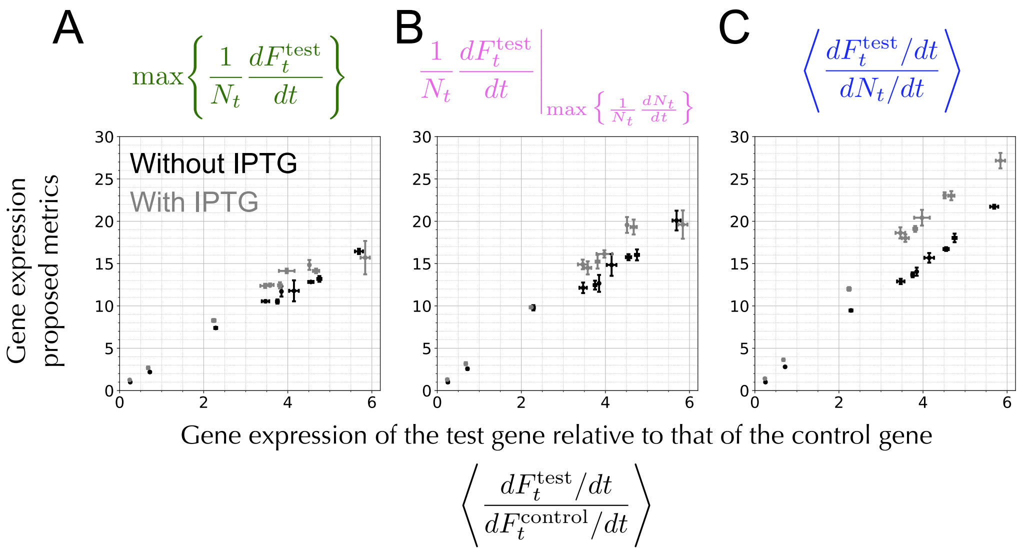

Figure S14: Transcription rate inferred from different methods for the same promoters as in Fig. 1D of the main text in absence or in presence of IPTG, corresponding to a closed or open loop. The results are essentially proportional to each other, with negligible differences induced by IPTG compared to differences due to promoters. The precision at which these differences are measured are, however, different from different methods (Fig. S15). We relie on the method presented in C for Fig. 1D of the main text.

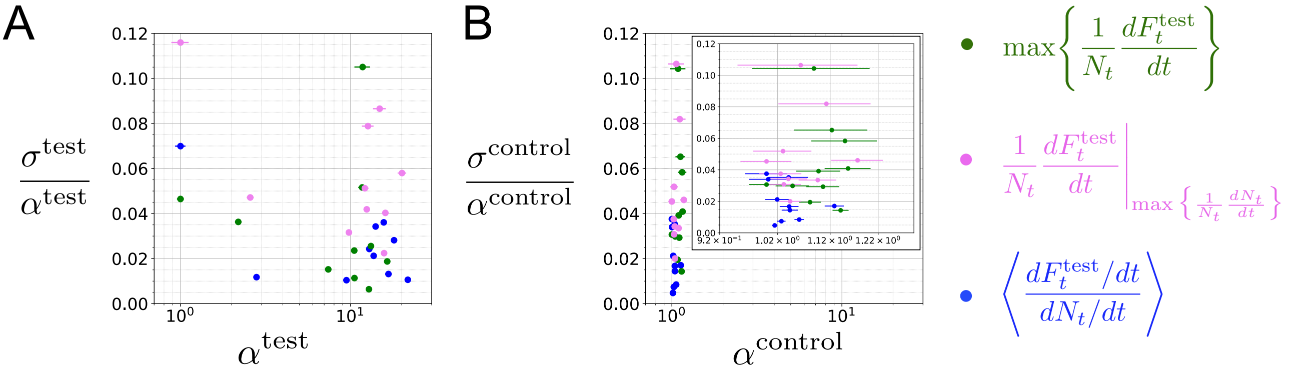

Figure S15: Precision of the different approaches for the data shown in Fig. 1 of the main text. Here we show statistics of standard deviations over replicates normalized by their associated average (Fano factor), both for the test gene (A), which spans a large dynamic range of promoter activities (Fig. S14) and for the control gene (B), which spans a short dynamic range of promoter activities. Error bars are standard deviations of Fano factors. The approaches show critical differences in their precision for the control gene which is subject to subtle changes, in which case our approach based on $(dF/dt)/(dN/dt)$ (in blue) leads to more precise estimations.

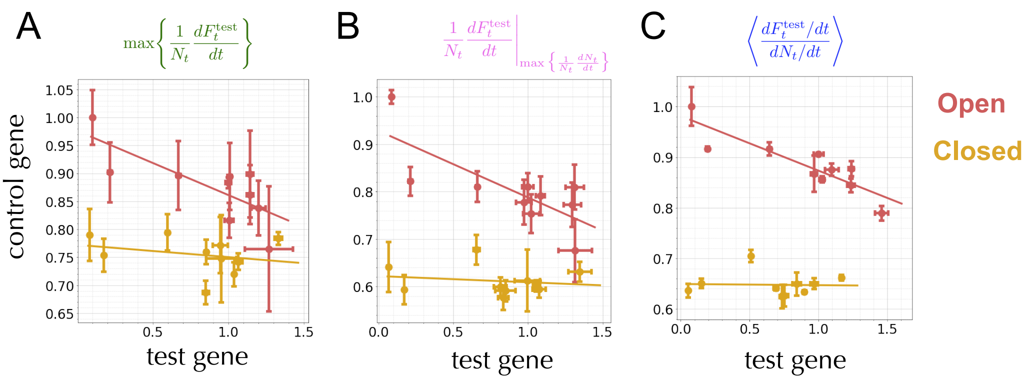

Figure S16: Robustness of the results of Fig. 1D to the definition of the expression rate – The qualitative result of lacI-based insulation are preserved, regardless of the inference method. Fig. 1D of the main text corresponds to panel C while panels A and B show the results obtained from the different approaches. The results are consistent but, as also shown in Fig. S15, our approach permits a more precise estimation of transcription rates.

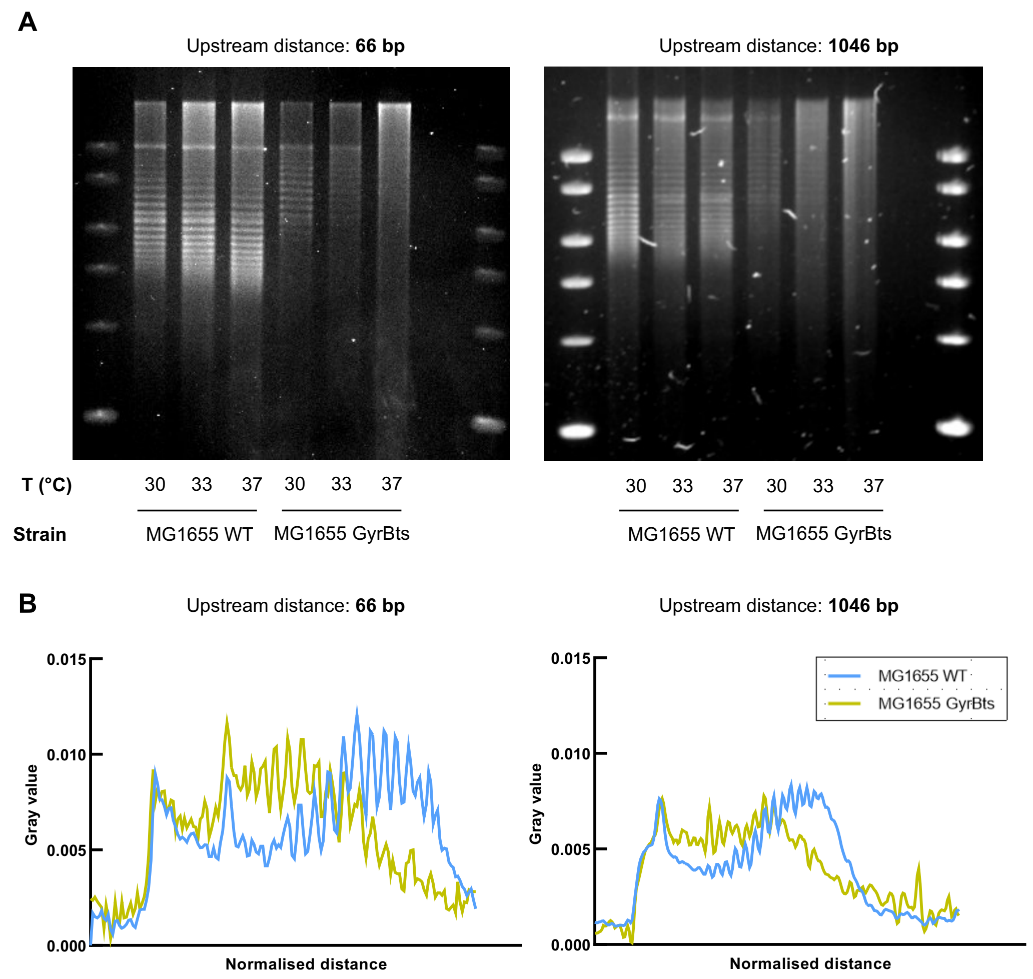

Figure S17: Effect of GyrB inactivation on plasmid supercoiling – **A.** One-dimensional chloroquine-agarose gel electrophoresis of plasmid LSOA (left panel, upstream distance of 66 bp) and LSOJ (right panel, upstream distance of 1046 bp). Plasmids were extracted from exponential phase for *E. coli* cells cultured in complemented m9 medium at 30, 33 or 37°C and analyzed on 0,8% agarose gel supplemented with 2.5 $\mu$g/mL of chloroquine. The gel electrophoresis was run in Tris-Borate-EDTA buffer containing 2.5 $\mu$g/mL chloroquine during 15 hours at 25V and stained by SYBR Green. **B.** Intensity graphs corresponding to the electrophoresis gels presented above. The migration profiles were obtained by using the plot function of ImageJ software on lines corresponding to the migration of plasmids extracted from cultures grown at 30°C. These results show that GyrB inactivation affects supercoiling.

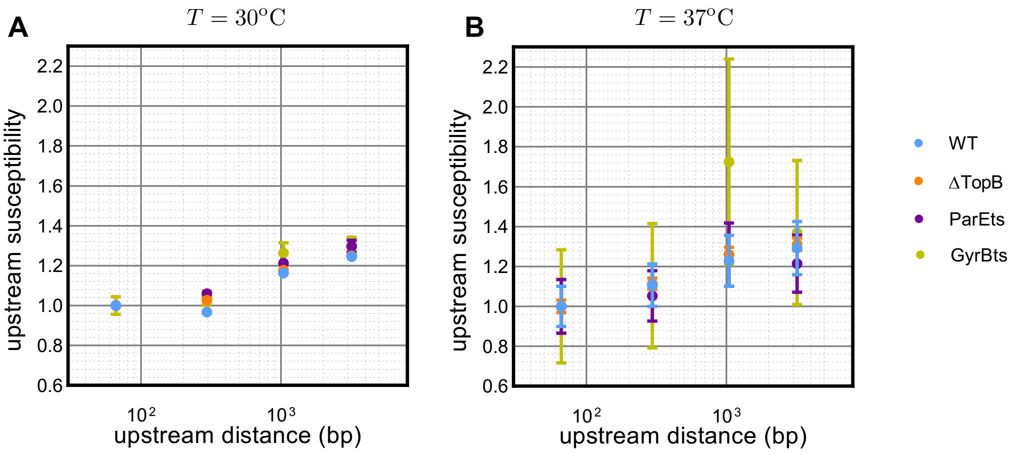

Figure S18: Susceptibility of gene expression to upstream context in WT or mutated *E. coli* MG1655 with deleted or inactivated topoisomerases or gyrase – The upstream susceptibility is defined as the ratio of the expression rate of the insulated gene with a long (∼ 3200 bp) distance to the upstream barrier over its expression rate with a short (∼ 100 bp) distance. Measurements were performed at 30°C (A) or 37°C (B) on WT, ΔTopB, ParEts and GyrBts *E. coli* strains (MG1655) where ΔTopB, ParEts and GyrBts have respectively TopoIII, TopoIV and gyrase inactivated. The results show no significant effect.

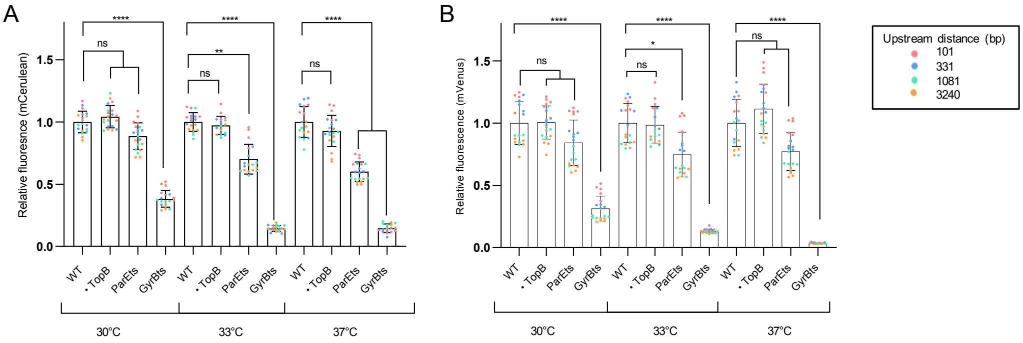

Figure S19: Effect of TopB deletion and inactivation of GyrB and ParE on reporter genes fluorescence measurements at different growth temperatures – Each point represents an individual well. The fluorescence values were normalized by the ones obtained with the WT strain. For each temperature, mutants are compared to WT by performing Kruskal-Wallis test. Promoter p01 (Table S1) is used for mCerulean. The results show that TopB deletion has no significant effect while inactivation of GyrB and ParE both lead to lower expression.

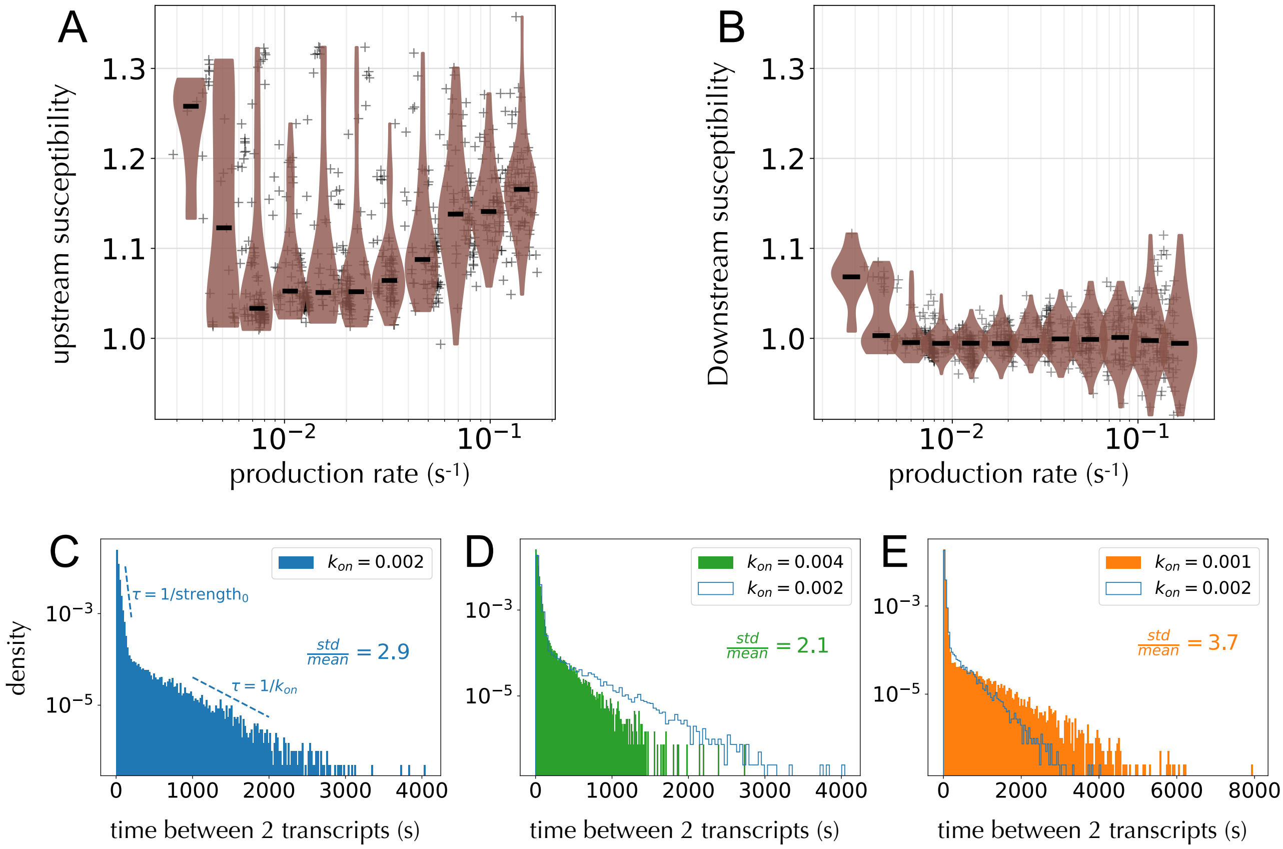

Figure S20: Motivated by previous *in vitro* and *in vivo* experiments [5], we tested whether adding a slow modulation of gyrase activity could reproduce the phenomenology of transcriptional bursting in addition to the upstream and downstream susceptibilities shown by our experimental results (Fig. 2 of main text). To this end, we assumed an additional process that modulates gyrase binding on a long temporal scale, using binding rate ($k_{\text{on}}$) and unbinding rate ($k_{\text{off}}$) on the order of $0.1\,\text{min}^{-1}$, while keeping the other parameters identical. In this new model, the effective rate of gyrase therefore changes with time, being equal to $\Lambda_s^{gyr} = -2.5$ Lk.s$^{-1}$ in the bound state and 0 in the unbound state. The time scale associated with $k_{\text{on}}$ and $k_{\text{off}}$ ($\sim 10\,\text{min}$) follows previously reported values for the slow modulation of gene expression in bacteria [6, 7]. Comparing panels A and B to Figure 5A, we find that this modification of the model does not change qualitatively the results for the upstream and downstream susceptibilities, except for very weak promoters where the time scale of the slow modulation starts to be on the order of the promoter production rate. Statistics is overall noisier than in the original model because of the long time scales – the plots here were produced by generating $10^5$ transcripts. In panels C, D, E, we show the distribution of times separating the production of two transcripts for the weak promoter in Fig. 5B (green curve and points) of the main text, for $k_{\text{on}} = k_{\text{off}} = 0.002\,\text{s}^{-1}$ (panel A), for $k_{\text{on}} = 0.004\,\text{s}^{-1}$ and $k_{\text{off}} = 0.002\,\text{s}^{-1}$ (panel B), hence mimicking an over-expression of gyrase with respect to panel A, and for $k_{\text{on}} = 0.001\,\text{s}^{-1}$ and $k_{\text{off}} = 0.002\,\text{s}^{-1}$ (panel C), hence mimicking an under-expression of gyrase. The three distributions reflect a non-Poissonian process characterized by a double-exponential decay with two distinct time scales: one associated with the original promoter strength $1/\text{strength}_0$ (i.e., without the slow alternation of gyrase activity) and a longer one given by $1/k_{\text{on}}$. We then report the value of the standard deviation over the mean, std/mean, for each plot, which is the counterpart of the Fano factor when considering only the production process; in particular, std/mean $= 1$ for an exponential distribution of times (Poissonian process). As reported in Ref. [5], we observe a less (more) bursty process for an over-expression (under-expression) of gyrase, which is characterized by a smaller (larger) value of std/mean. The present extension of the model is not intended to be physically realistic but illustrates how our results and interpretations are consistent with the observation of transcriptional bursts. A more realistic model will be the subject of future work.
